# Regulation with cell size ensures mitochondrial DNA homeostasis during cell growth

**DOI:** 10.1101/2021.12.03.471050

**Authors:** Anika Seel, Francesco Padovani, Alissa Finster, Moritz Mayer, Daniela Bureik, Christof Osman, Till Klecker, Kurt M. Schmoller

## Abstract

To maintain stable DNA concentrations, proliferating cells need to coordinate DNA replication with cell growth. For nuclear DNA, eukaryotic cells achieve this by coupling DNA replication to cell cycle progression, ensuring that DNA is doubled exactly once per cell cycle. By contrast, mitochondrial DNA replication is typically not strictly coupled to the cell cycle, leaving the open question of how cells maintain the correct amount of mitochondrial DNA during cell growth. Here, we show that in budding yeast, mitochondrial DNA copy number increases with cell volume, both in asynchronously cycling populations and during G1 arrest. Our findings suggest that cell-volume-dependent mitochondrial DNA maintenance is achieved through nuclear encoded limiting factors, including the mitochondrial DNA polymerase Mip1 and the packaging factor Abf2, whose amount increases in proportion to cell volume. By directly linking mitochondrial DNA maintenance to nuclear protein synthesis, and thus cell growth, constant mitochondrial DNA concentrations can be robustly maintained without a need for cell-cycle-dependent regulation.

## Introduction

As cells grow during the cell cycle, they need to double their DNA content so that each daughter cell obtains the appropriate amount. In fact, it is one of the major tasks of the eukaryotic cell cycle to ensure that nuclear DNA is replicated once – and only once – during S-phase, and decades of extensive research provided us with a detailed molecular understanding of this process (Sclafani and Holzen, 2007; Ekundayo and Bleichert, 2019). By contrast, how this is achieved for mitochondrial DNA (mtDNA), the other major type of DNA in virtually all eukaryotes, is largely unclear.

In many organisms, including humans and yeasts, mtDNA encodes proteins essential for oxidative phosphorylation and is typically present in many copies (Gustafsson et al., 2016; Aretz et al., 2020). mtDNA is organized in ‘nucleoids’, nucleoprotein complexes that are distributed throughout the mitochondrial network and can contain one or several copies of mtDNA (Kukat et al., 2011; Göke et al., 2020). While several regulators of mtDNA that affect mtDNA copy number have been identified (Göke et al., 2020), including the nucleoid protein TFAM (Ekstrand et al., 2004) and its homolog Abf2 in yeast (Zelenaya-Troitskaya et al., 1998), mtDNA polymerase (Stumpf et al., 2010), and helicases (Tyynismaa et al., 2004; Taylor et al., 2005), how cells maintain the correct number of mtDNA throughout cell growth is unknown. In contrast to replication of nuclear DNA, mtDNA replication is not strictly coupled to cell cycle progression. While some studies report cell-cycle-dependent modulation of mtDNA replication rates for human cells (Lee et al., 2007; Chatre and Ricchetti, 2013; Sasaki et al., 2017), mtDNA replication occurs throughout the cell cycle and even continues during long cell cycle arrests (Petes and Fangman, 1973; Wells, 1974; Newton and Fangman, 1975; Conrad and Newlon, 1982). However, if mtDNA replication is not controlled by cell cycle progression, how can cells then coordinate the amount of mtDNA produced with cell growth?

One alternative to regulation by the cell cycle is that mitochondrial homeostasis is directly linked to cell size. Indeed, it has been shown that the total amount of mitochondria in budding yeast (Rafelski et al., 2012), HeLa (Posakony et al., 1977), mouse liver (Miettinen et al., 2014), Jurkat and *Drosophila* Kc167 cells (Miettinen and Björklund, 2016) increases roughly in proportion to cell volume. In addition, the number of nucleoids in budding yeast correlates with mitochondrial network volume (Osman et al., 2015), and nucleoid number in fission yeast increases with increasing cell volume (Jajoo et al., 2016) – suggesting that also mtDNA copy number might be linked to cell volume. However, direct evidence for a role of cell volume for mtDNA homeostasis is missing.

Here, we show that in budding yeast, the number of mtDNA copies and nucleoids increases in direct proportion to cell volume. We find that mtDNA maintenance is limited by nuclear encoded proteins whose abundance increases with cell volume. Supported by mathematical modelling, our results suggest a mechanism in which the overall increase of cellular protein synthesis with increasing cell volume couples mtDNA copy number to cell volume, achieving robust mtDNA homeostasis during cell growth and cell cycle progression.

## Results

### mtDNA copy number increases with cell volume

As a first step to understand the role of cell size in the regulation of mtDNA copy number, we measured the dependence of mtDNA copy number on cell volume in budding yeast. We used haploid and diploid strains carrying the cell size regulator *WHI5* under control of a synthetic, β-estradiol inducible promoter (Ottoz et al., 2014). Whi5 modulates G1 duration by inhibiting the major G1/S transcription factor SBF. Overexpression of Whi5 by addition of β-estradiol to the media therefore initially causes a prolonged G1 phase and an increase of cell volume. Eventually, cells reach a cell volume large enough to allow division even at the increased Whi5 expression (Schmoller et al., 2015), and after 24 hours of growth in exponential phase, this results in a new steady state of asynchronous cell populations with increased mean cell volumes (Claude et al., 2021; Kukhtevich et al., 2020) (Fig. 1a). We grew cells on synthetic complete media with 2% glycerol and 1% ethanol as non-fermentable carbon source (SCGE). Using β-estradiol concentrations ranging from 0 to 30 nM (haploid) or 60 nM (diploid) as well as wild-type strains without inducible-Whi5, we obtained steady-state cultures with a more than 4-fold range in mean cell volume (Supplementary Fig. 1), but similar doubling times and only moderately shifted cell cycle fractions (Claude et al., 2021). For each culture, we measured cell volume using a Coulter counter, determined bud fractions by visual inspection, purified DNA, and performed qPCR measurements on nuclear and mtDNA to determine the average number of mtDNA copies per cell. For both haploids and diploids, we found that mtDNA copy number increases roughly in direct proportion with cell volume (Fig. 1b).

**Figure 1.**
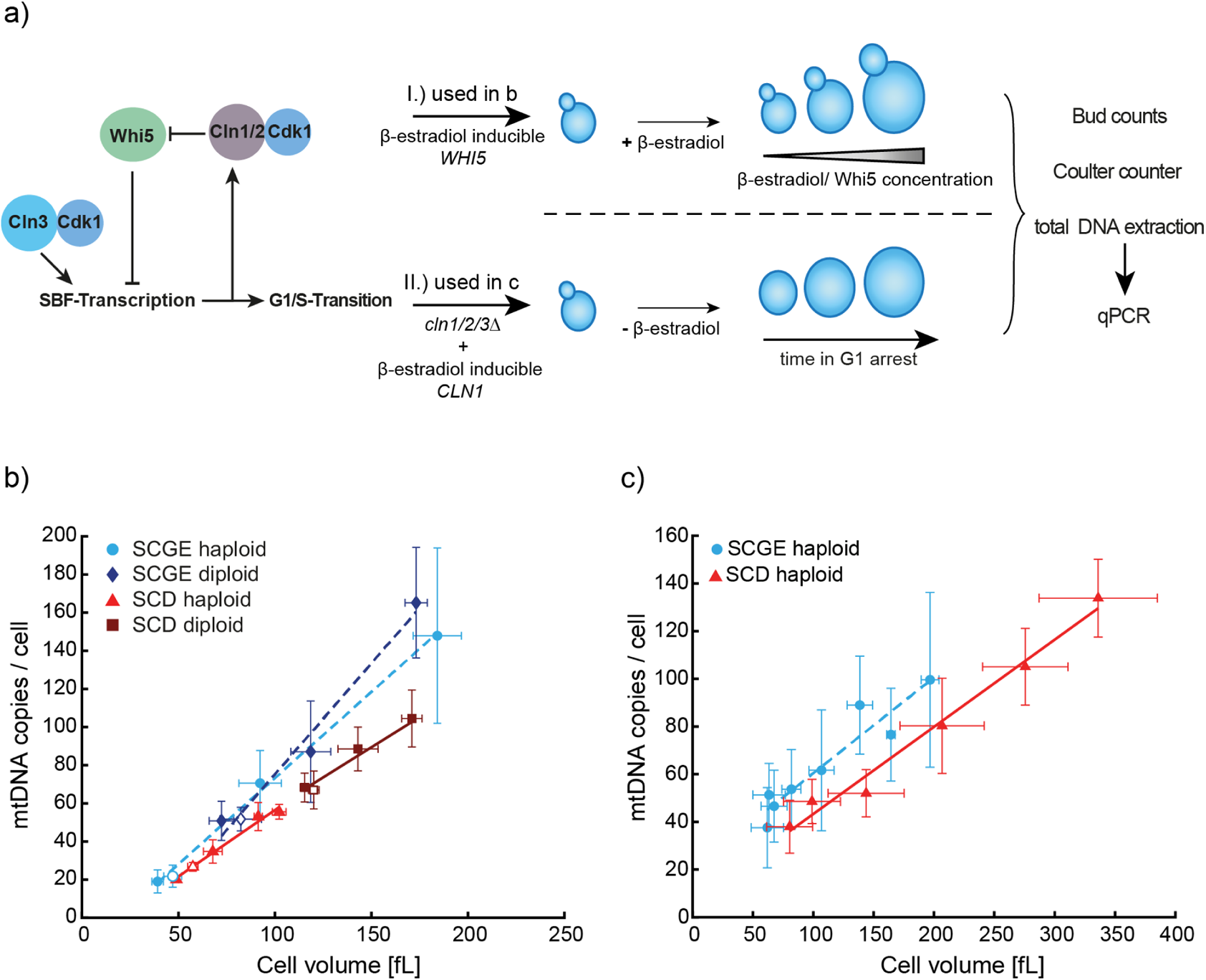
mtDNA increases with cell volume and is modulated by nutrients. **a)** Cell volume was manipulated using two different genetic approaches. I.) Whi5 concentration was controlled by a β-estradiol-inducible promoter. Addition of higher β-estradiol concentrations leads to a prolonged G1-phase resulting in bigger cell volumes in asynchronous steady state populations. II.) A *cln1/2/3* deletion strain with Cln1 expressed from a β-estradiol-inducible promoter was used (Ewald et al., 2016). In this strain, β-estradiol is necessary for cell proliferation and removal leads to a G1-arrest and continuously increasing cell volumes. **b)** mtDNA copy number as a function of cell volume in asynchronous steady state populations. Haploid and diploid wild-type (open symbols) and Whi5-inducible strains (filled symbols) in the absence or presence of different β-estradiol concentrations were grown on SCGE (dashed line) or SCD medium (solid line). After total DNA extraction, mtDNA copy number was determined by measuring the relative concentration of mtDNA and nuclear DNA. mtDNA concentration was normalized on nuclear DNA concentration and the budding index was used to estimate nuclear DNA copies per cell. Mean cell volumes of cell populations were measured with a Coulter counter. **c)** mtDNA copy numbers as a function of cell volume during G1 arrest. Cells were arrested in G1 and harvested every hour for 6 hours (SCD) or 8 hours (SCGE), starting directly after β-estradiol removal. mtDNA copy numbers were measured with qPCR, cell volumes were measured with a Coulter counter. Lines show linear fits to the means. Error bars indicate standard deviations of at least 3 replicates (**b, c**).

We next asked whether this increase of mtDNA amount with cell volume is specific to non-fermentable media, in which respiratory activity and thus also functional mtDNA is essential. We therefore repeated the experiments using synthetic complete media with 2% glucose (SCD), in which mtDNA is not essential for budding yeast growth. Again, we found that the amount of mtDNA increases with cell volume, but at a given cell volume, cells grown on SCD have less mtDNA compared to cells grown on SCGE (Fig. 1b).

To test whether the observed increase of mtDNA is due to the increase in cell volume rather than linked to Whi5 overexpression, we then sought for an alternative approach to obtain cells of different volume. We used a haploid strain in which all three endogenous G1-cyclins (*CLN1/2/3*) were deleted, and which is kept alive by a β-estradiol-inducible copy of *CLN1* (Ewald et al., 2016). Upon removal of β-estradiol from the media, the cells arrest in G1 while continuously growing (Supplementary Fig. 2). We collected cells at different time points of the G1 arrest, again measuring cell volume, bud fractions, and mtDNA copy number (Fig. 1a). Consistent with mtDNA being replicated also in G1 (Petes and Fangman, 1973; Newton and Fangman, 1975), and confirming the results obtained with the Whi5-inducible system, we found that as cells grow during G1, mtDNA copy number continuously increases (Fig. 1c). Again, we found that mtDNA copy number is lower on SCD than on SCGE media.

### Nucleoid number increases with cell volume

mtDNA is organized in nucleoprotein complexes called nucleoids, which in budding yeast are distributed throughout the mitochondrial network (Osman et al., 2015). In principle, the increase in mtDNA with cell volume could either be due to an increased number of nucleoids, or an increase in the number of mtDNA copies per nucleoid. To distinguish between the two scenarios, we adapted a previously established system to visualize mtDNA in live cells (Osman et al., 2015). Briefly, we introduced inducible-Whi5 into haploid and diploid strains in which LacO arrays have been stably integrated into the mitochondrial genome (Fig. 2a). A constitutively expressed LacI tagged with two copies of mNeon and a mitochondrial targeting sequence then binds to the LacO arrays, resulting in fluorescent foci detectable with confocal microscopy (Fig. 2b; Supplementary Fig. 3a). At the same time, we used the fluorescent protein mKate2 targeted to the mitochondrial matrix to visualize the mitochondrial network. To maximize the observable range of cell volumes, we induced Whi5 expression in cells grown either on SCGE or SCD media with different concentrations of β-estradiol and then imaged cells with 3D-confocal microscopy. Using a custom image analysis pipeline, we then segmented cells based on bright field images, and segmented the mitochondrial network and identified mtDNA foci in 3 dimensions from fluorescence signals. Consistent with a previous report (Rafelski et al., 2012), the volume of the mitochondrial network increases in proportion with cell volume, with cells on SCD having less mitochondrial network than cells on SCGE (Fig. 2c). At the same time, we found that also the number of mtDNA foci (which we interpret as nucleoids) increases with cell volume, indicating an increase in the number of nucleoids rather than in the mtDNA copy numbers per nucleoid. In line with the reduced number of mtDNA copies per cell (Fig. 1b-c), we detected fewer nucleoids at a given cell volume when cells were grown on SCD (Fig. 2d).

**Figure 2.**
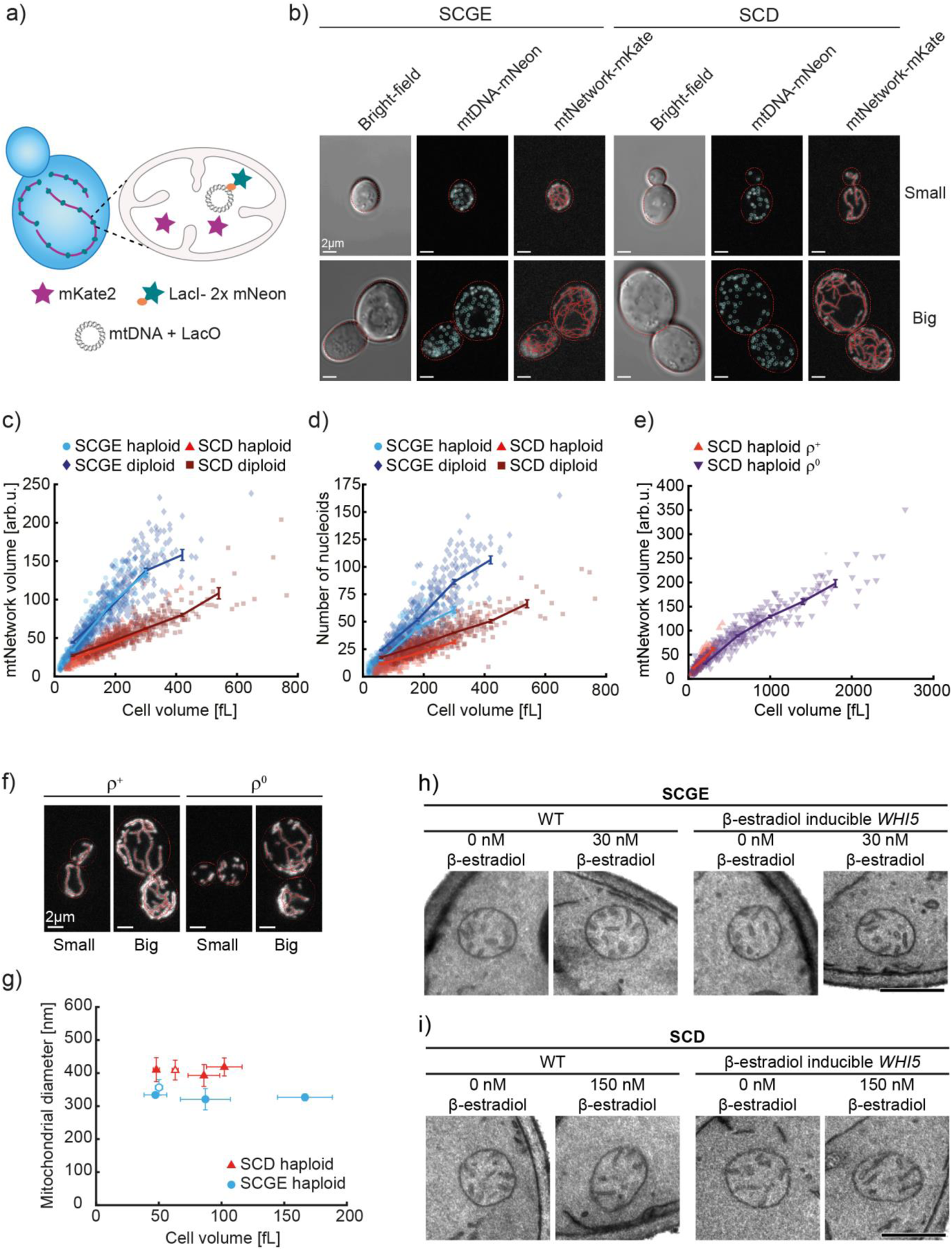
Number of nucleoids and mitochondrial network volume increase with cell volume. **a)** For live-cell imaging, strains with LacO-repeats integrated into mtDNA were used (Osman et al. 2015). LacI-2xmNeon was expressed from nuclear DNA and targeted to mitochondrial matrix where it binds LacO-repeats. By expressing mitochondrially targeted mKate2, the mitochondrial matrix was visualized. **b)** Representative bright-field and confocal live-cell images (maximum intensity projections) of Whi5-inducible diploid cells without (small) or with 60 nM (SCGE) / 150 nM (SCD) β-estradiol (big) are shown together with cell and mitochondrial network segmentations as well as identified mtDNA foci. Note that the skeletonization of the segmentation mitochondrial network was only used here for visual representation. Corresponding images without network segmentation and nucleoid detection are shown in Supplementary Fig. 3a. **c)** Mitochondrial network volume as a function of cell volume for Whi5-inducible cells grown on SCGE and SCD medium with different β-estradiol concentrations. **d)** Number of nucleoids per cell as a function of cell volume, for the same cells as in c. **e)** The increase of mitochondrial network volume with cell volume does not depend on mtDNA. *MIP1* was deleted in a Whi5-inducible haploid strain to generate a ρ^0^ strain. Mitochondrial network volume is shown as a function of cell volume and compared to ρ^+^ cells of the parental strain (data from 2d). Cells were grown on SCD. **b-e)** Image analysis was performed in 3D. Per condition, 3 biological replicates with n=50 images each were analyzed (total n=150). Lines connect binned means (shown at the center of the respective bin) with error bars indicating standard errors. **f)** Representative confocal live-cell images (maximum intensity projections) of ρ^+^ and ρ^0^ cells without (small) or with 150 nM β-estradiol (big). Corresponding images without network segmentation are shown in Supplementary Fig. 3b. **g-i)** Haploid wild-type (open symbols) and Whi5-inducible cells (filled symbols) were grown in SCD or SCGE containing different concentrations of β-estradiol. The cells were chemically fixed and analyzed by transmission electron microscopy. **g)** Mitochondrial diameter as a function of cell volume. For each sample, the diameter of 100 mitochondria was measured from electron micrographs and cell volumes were calculated from DIC images taken of the same cultures that were used for chemical fixation. Shown is the mean of the means from three experiments, error bars indicate standard deviations of the means. Cells grown in SCD and SCGE were analyzed in separate experiments. See also Supplementary Fig. 4a-d. **h-i)** Representative electron micrographs of mitochondria of wild-type (WT) and Whi5-inducible cells grown either in SCGE (h) or SCD (i) medium containing the indicated concentrations of β-estradiol. Scale bars represent 500 nm. Additional images are shown in Supplementary Fig. 4e-f.

### The increase of mitochondrial network with cell volume is independent of mtDNA

So far, we have shown that larger cells not only have more mitochondrial network, but also an increased amount of mtDNA and nucleoids. We therefore wondered whether the increase in mitochondrial network volume is a causal consequence of the increased mtDNA copy number. To address this question, we deleted the only mitochondrial DNA polymerase in yeast, *MIP1*, which results in loss of mtDNA (Supplementary Table 1). Since mtDNA is not essential for budding yeast to grow on fermentable media, we could grow this strain on SCD media, and measure mitochondrial network volume as a function of cell volume. We noticed that these cells without mitochondrial DNA, called ρ^0^ cells, show a larger variability of cell volume, in particular after Whi5 over-expression, and exhibit altered network morphology (Supplementary Fig. 3b-c). Nevertheless, the network volume still increases with cell volume, and at a given cell volume is similar to that in wild-type cells (Fig. 2e-f). Thus, the regulation of mitochondrial network volume with cell volume occurs either upstream or independent of mtDNA.

### Mitochondrial diameter is independent of cell volume

We observe that mitochondrial network volume, mtDNA amount, as well as nucleoid number all increase with cell volume. Together with previous reports on yeast (Rafelski et al., 2012) and mammalian cells (Posakony et al., 1977; Miettinen et al., 2014), our results suggest that the total amount of mitochondria increases in larger cells but that the local mitochondrial structure is rather constant. To test if mitochondrial diameter changes with increasing cell volume, we analyzed haploid wild-type and Whi5-inducible cells grown in SCD or SCGE media in the absence or presence of β-estradiol by transmission electron microscopy and measured mitochondrial width. We observed no changes in mean mitochondrial diameter associated with increasing cell volume upon Whi5 overexpression, irrespective of whether the cells were grown in SCD or SCGE (Fig. 2g; Supplementary Fig. 4a-d). Furthermore, we observed no obvious cell-volume-dependent alterations of mitochondrial inner membrane structure (Fig. 2h-i, Supplementary Fig. 4e-f). Taken together, our results indicate that the mitochondrial diameter is not influenced by cell volume in budding yeast.

### Amount of mtDNA maintenance factors increases with cell volume

In summary, we have shown that mtDNA copy number is tightly linked to cell volume both in asynchronous as well as G1-arrested cells. In principle, this coupling of mtDNA to cell volume allows cells to maintain mtDNA concentrations during cell growth: If mtDNA copy numbers are set by cell volume at any point during the cell cycle, constant concentrations can be achieved without any dedicated regulation of mtDNA replication with cell cycle progression. This then raises the question of how mtDNA copy number can be coordinated with cell volume.

Most proteins increase in abundance as cells get bigger, maintaining constant concentrations (Swaffer et al., 2021a). This overall increase of protein production is thought to be due to an increased abundance of ribosomes (Marguerat and Bähler, 2012; Schmoller and Skotheim, 2015), but also an increase of global transcription, and thus mRNA amounts (Wu et al., 2010; Zhurinsky et al., 2010; Padovan-Merhar et al., 2015; Sun et al., 2020; Swaffer et al., 2021b).

One possible mechanism of how mtDNA number could be linked to cell volume is that similar to most genes, mitochondrial maintenance factors encoded by the nuclear genome might be higher expressed in larger cells. This could then lead to higher protein amounts, potentially also including the proteins limiting for mtDNA maintenance. We therefore asked whether the expression of factors known to be necessary for mtDNA maintenance indeed increases with cell volume.

As a first step to address this question, we determined the dependence of transcript concentration on cell volume for several nuclear encoded mitochondrial factors. To this end, we re-analyzed by RT-qPCR RNA samples of a Whi5-inducible strain grown on SCGE that we recently used to determine the concentration of histone transcripts as a function of cell volume (Claude et al., 2021). As we have shown previously, the transcript concentration of the control-gene *ACT1* is maintained nearly constant with cell volume. In contrast, the concentration of histone mRNA decreases in inverse proportion with cell volume to maintain constant histone amounts (Claude et al., 2021). Here, we find that the transcript concentration of all nuclear encoded mitochondrial factors analyzed only slightly decreases with cell volume, resulting in significantly increased transcript amounts in large cells (Fig. 3a). To validate this finding, we could make use of two previously published datasets (Swaffer et al., 2021a) that measured the cell-size dependence of transcripts 1.) in budded cells sorted by cell size (total protein content) using flow cytometry, and 2.) during the first cell cycle of cells released from G1 arrests of varying time, and thus of cells with different volumes. Again, we find that similar to most transcripts, including those of RNA Polymerase II subunits and *ACT1*, the transcripts of factors involved in mtDNA maintenance are kept at largely cell-size independent concentrations (Fig. 3b).

**Figure 3.**
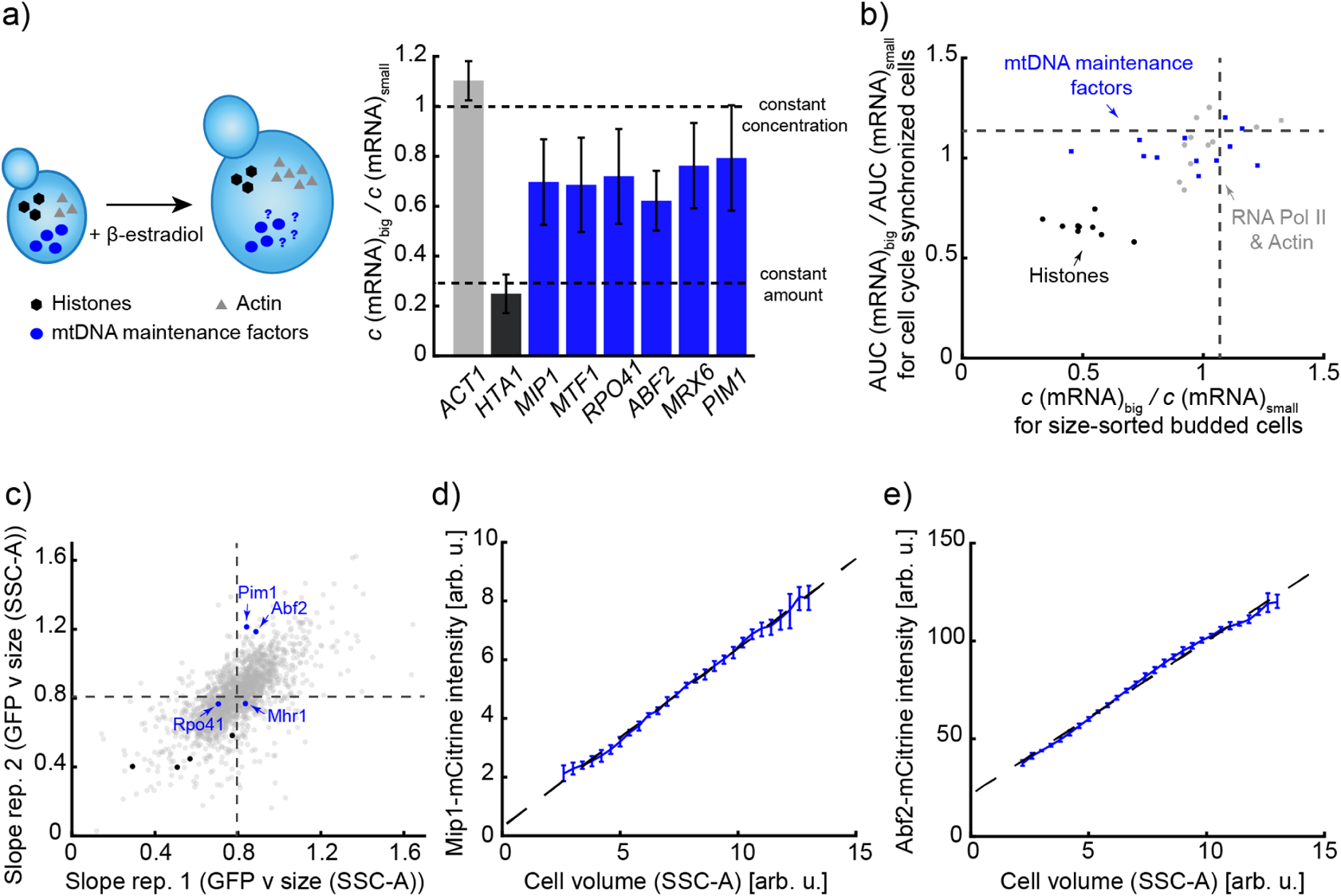
Amount of nuclear encoded mtDNA maintenance factors increases with cell volume. **a)** mRNA amounts of nuclear encoded mitochondrial proteins increase with cell volume. Cells were grown on SCGE with 0 (small) or 30 nM β-estradiol (big), followed by total RNA extraction and RT-qPCR. mRNA concentrations were normalized on *RDN18* and the ratio of concentrations in big (30 nM) and small (0 nM) haploid cell populations were calculated. Bars indicate mean of 3 replicates with error bars indicating standard deviations. Cell volumes were measured with a Coulter counter to estimate the concentration ratio expected for an mRNA that is maintained at constant amount. *MIP1*, *MTF1*, *RPO41*, *ABF2*, *MRX6* and *PIM1* data were generated from RNA samples from Claude et al., 2021. *ACT1* and *HTA1* measurements were directly taken from Claude et al., 2021. **b)** Concentration of nuclear transcripts encoding for proteins involved in mitochondrial mtDNA maintenance are largely constant with cell volume in two transcriptomics data-sets from (Swaffer et al., 2021a): The ratios in big and small cells of the relative expression in size-sorted budded cells, and of the Area Under the Curve (AUC) of the relative expression during cell cycle progression after different durations of G1 arrest are shown for mitochondrial proteins (blue), control genes (RNA polymerase II and *ACT1*) known to scale with cell volume (grey), and histones (black), whose concentration decreases in big cells. Medians of all transcripts are shown as dashed lines. **c)** Analysis of the cell-volume-dependence of GFP-fusion proteins performed by Swaffer et al. based on data by Parts et al. (Parts et al., 2014). The normalized slope of a linear fit to the GFP intensity as a function of cell size (SSC-A) for budded cells in 2 biological replicates was used to estimate the cell-volume-dependence for each fusion protein. Similar to most proteins (grey), the amount of mtDNA maintenance factors (blue) increases roughly in direct proportion with cell volume. In contrast, the concentration of histones (black) decreases in big cells. Mean slopes for all fusion proteins included in the data set are shown as dashed lines. **d-e)** Flow-cytometry was used to measure total cellular mCitrine fluorescence intensity in haploid strains in which either *MIP1* (d) or *ABF2* (e) were endogenously tagged. SSC-A-signal was used as a measurement for cell volume (Supplementary Fig. 6). Binned means after background correction using a non-fluorescent strain are shown with estimated experimental errors (see methods for details). Dashed lines show linear fits to the binned means.

As a next step, we asked whether the increasing transcript amounts indeed lead to larger amounts of the corresponding proteins. Again, we could make use of an analysis performed by Swaffer et al. (Swaffer et al., 2021a): using previously published flow-cytometry data on a collection of strains in which each open reading frame (where possible) was tagged with GFP (Parts et al., 2014), the dependence of protein amounts on cell size was analyzed. While many of the mtDNA maintenance factors were excluded from this data-set for technical reasons, in particular due to low expression levels, all mtDNA factors included (Abf2, Mhr1, Pim1, Rpo41) showed an increase of protein amount with cell size similar or stronger than the average of all measured proteins (Fig. 3c).

To further confirm that the amount of proteins necessary for mtDNA maintenance increases with cell volume, we constructed haploid strains in which we endogenously tagged the mtDNA polymerase *MIP1* as well as the mtDNA packaging factor *ABF2* (the budding yeast TFAM protein), respectively, with the fluorescent protein *mCitrine*. We ensured that the tagged proteins are functional by testing growth on SCGE and measuring mtDNA concentrations with qPCR (Supplementary Fig. 5). In the case of Mip1-mCitrine, we found that the strain with the tagged allele has increased amounts of mtDNA (Supplementary Fig. 5a), but still shows the typical increase of mtDNA copy number with cell volume. Thus, the mechanism ensuring the cell-volume-dependence is still intact, suggesting that also the regulation of Mip1 with cell volume is not dramatically impaired by the tag (Supplementary Fig. 5e). Using mCitrine fluorescence intensity as a proxy for protein amount, and side scatter as a measure for cell volume (Supplementary Fig. 6), we then determined the cell-volume-dependence of Mip1 and Abf2 protein amounts with flow cytometry. Consistent with the transcript measurements, we found that the amounts of both Mip1-mCitrine and Abf2-mCitrine strongly increase with cell volume (Fig. 3d-e).

### Mip1 and Abf2 are limiting for mtDNA maintenance

Our results suggest that in larger cells, the proteins required for mtDNA replication and maintenance are present at higher numbers. If those proteins are limiting for mtDNA maintenance, meaning that a change in the protein abundance would cause a proportional change in the mtDNA copy number, this cell-volume-dependent increase in proteins could explain why larger cells have more mtDNA. To test if this is the case, and if so, which proteins are limiting, we created a series of hemizygous diploid strains, in each of which we deleted one allele of a gene involved in mtDNA maintenance (Contamine and Picard, 2000), including the mitochondrial DNA polymerase Mip1 (Stumpf et al., 2010), the DNA packaging protein Abf2 (Zelenaya-Troitskaya et al., 1998), the ssDNA-binding protein Rim1 (Van Dyck et al., 1992), helicases (Crider et al., 2012; Muellner and Schmidt, 2020; Sedman et al., 2000), and proteins involved in DNA recombination (Ling and Yoshida, 2020). Because most budding yeast genes do not exhibit dosage compensation at the transcript (Torres et al., 2016) or protein level (Springer et al., 2010), hemizygous diploids will in most cases show a 50% decrease of the corresponding transcript and protein. Should this protein then be perfectly limiting for mtDNA maintenance, we expect a 50% reduction of the mtDNA copy number (Fig. 4a).

**Figure 4.**
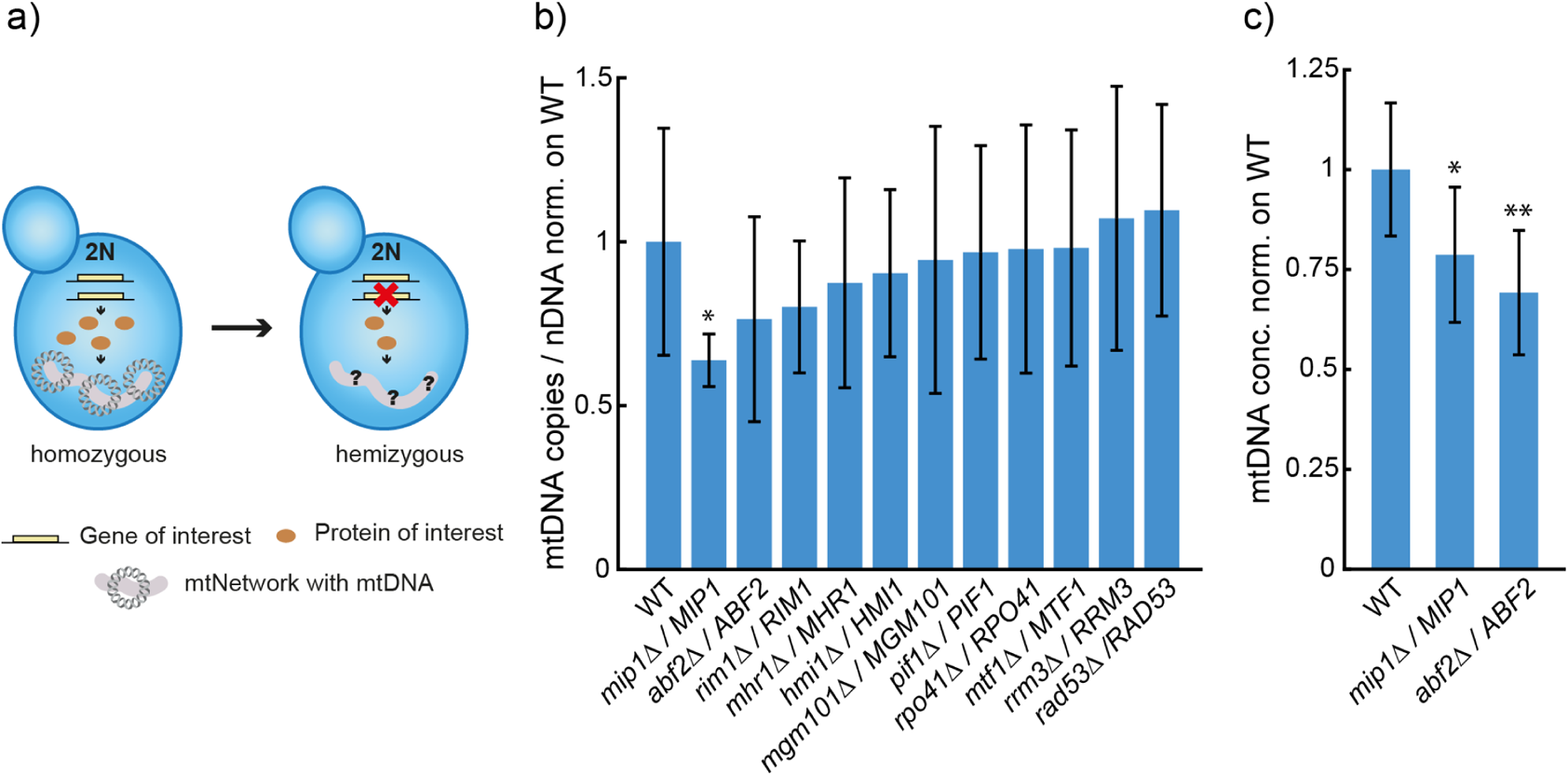
Hemizygous screen identifies limiting factors for mtDNA maintenance. **a)** To reduce the expression level of potentially limiting mtDNA maintenance factors, hemizygous diploid strains were constructed by deleting one allele of the gene of interest. **b)** Hemizygous diploid strains were grown on SCGE. mtDNA copy number per nuclear DNA as determined by DNA-qPCR were normalized on wild-type. Bars represent the mean of at least 3 replicates and error bars indicate standard errors. **c)** Independent validation of hemizygous *MIP1* and *ABF2* strains shown as mtDNA concentration (mtDNA copy number per cell / cell volume) normalized on WT. Significances were determined by a two-tailed, two-sample t test (* p<0.05, ** p<0.01).

For each hemizygous strain, we performed qPCR experiments to measure the ratio of mtDNA to nuclear DNA copies. We also measured cell volume, but did not find any major changes caused by the hemizygous deletions (which would cause changes in mtDNA copy number) (Supplementary Fig. 7a). As shown in Fig. 4b, we found that reducing the gene dosage of *MIP1* and *ABF2* had the strongest effect on mtDNA copy number. However, as validated by additional independent experiments (Fig. 4c), even diploids hemizygous for *MIP1* or *ABF2* showed only a reduction to 79% and 69%, respectively, of the mtDNA concentration. To rule out the possibility that we do not see a reduction of mtDNA to 50% because *MIP1* and *ABF2* exhibit dosage compensation, we measured the expression level. For both hemizygotes, we did not observe clear evidence for dosage compensation (Supplementary Fig. 7b-d) – consistent with a previous study that showed the absence of dosage compensation on the protein level in a strain heterozygous for fluorescently tagged Abf2 (Springer et al., 2010). Taken together, our results suggest that while mtDNA copy number is sensitive to the concentration of several proteins, none of the proteins we tested is perfectly limiting.

### Mathematical model explains coordination of mtDNA with cell volume

Since we found that the cell-volume-dependent increase of mtDNA cannot be trivially explained by a proportional increase of a single perfectly limiting factor, we decided to use mathematical modelling to obtain a better understanding of how several partially limiting components of the mtDNA maintenance machinery could contribute to mtDNA homeostasis. Because *MIP1* and *ABF2* hemizygotes showed the strongest reduction of mtDNA copy number, we decided to build a simple model focusing on the role of only these two proteins, neglecting the smaller contribution as limiting factors of other proteins.

In essence, and ignoring cell-to-cell variability, mtDNA copy number depends on the rates of mtDNA replication, degradation, and dilution by cell growth. While Mip1 might also affect mtDNA stability via its reported exonuclease activity (Viikov et al., 2012), we aimed to build a minimal model to understand the underlying principles, and therefore decided to only consider its obvious role in replication. Similarly, also Abf2 might be important for both replication and stability of mtDNA. If *abf2* deletion mutants are grown on fermentable media, they rapidly lose mtDNA. Nevertheless, *abf2Δ* cells can be grown on non-fermentable media, where they can maintain a pool of mtDNA over many generations (Diffley and Stillman, 1991), demonstrating that Abf2 is not essential for mtDNA replication. Instead, the compaction of mtDNA mediated by Abf2 seems to be important for mtDNA stability, suggesting that including this function of Abf2 in the model could be sufficient to understand the fundamental regulatory principles.

In the model, we account for these considerations by describing replication as a process that occurs at a rate that is determined by the concentrations of the mtDNA itself, *n*, as well as that of the mtDNA polymerase Mip1, *m*, with Michaelis-Menten-like kinetics 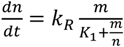 (Fig. 5a-b). Here, *k_R_* and *K_1_* are constants describing the maximal rate of replication per mtDNA and the dissociation constant of *Mip1* and mtDNA, respectively.

**Figure 5.**
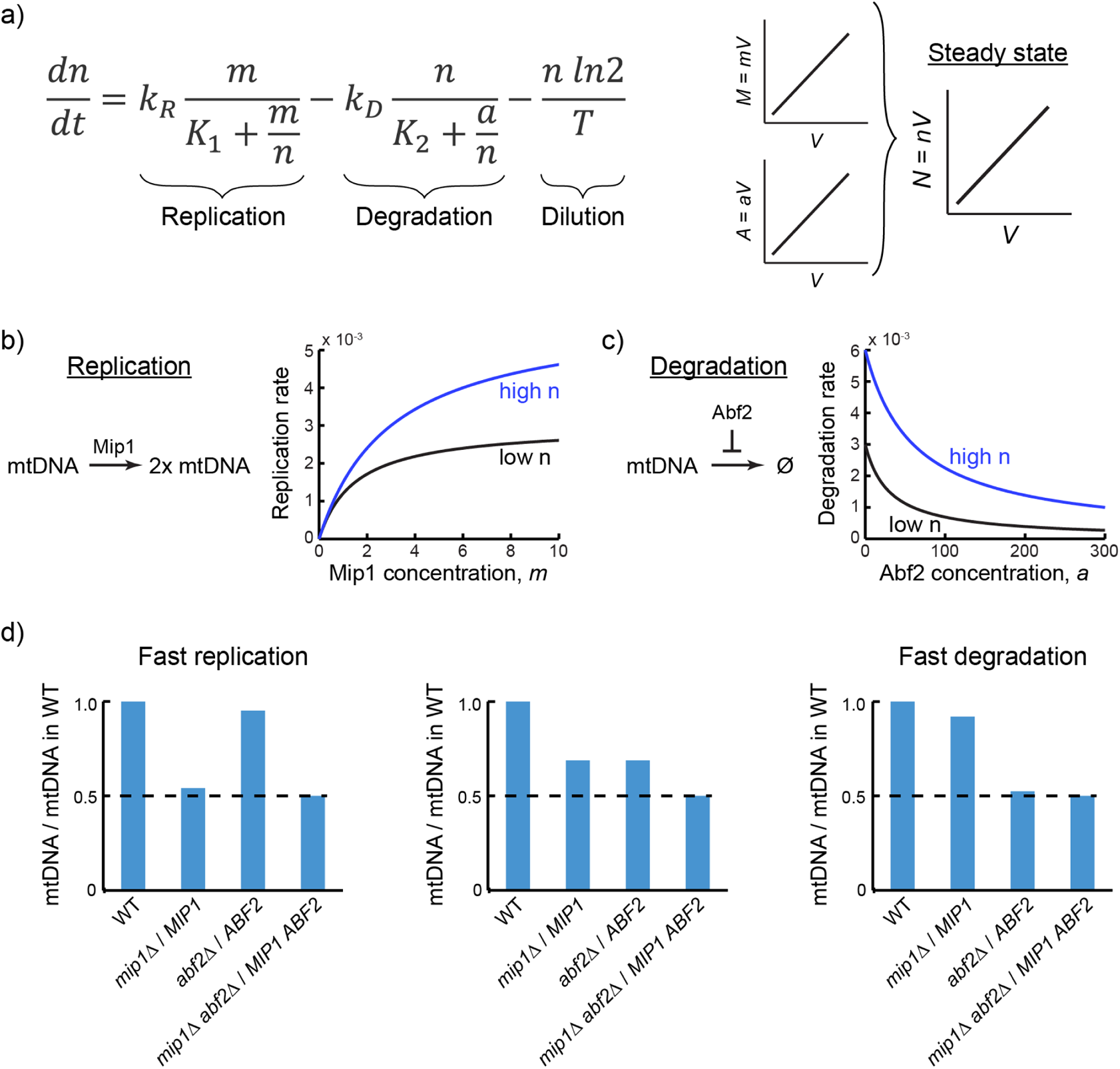
Mathematical model shows that limiting factors can couple mtDNA copy number to cell volume. **a)** The dynamic change of mtDNA concentration, *n*, is determined by mtDNA replication, degradation, and dilution due to cell growth. Cell growth is modeled as exponential. **b)** Replication is modeled as being limited by the mtDNA polymerase Mip1 at low Mip1 concentrations, *m*, and by *n* for saturating *m*. **c)** In the absence of Abf2, mtDNA degradation is modeled as an exponential decay. Increasing ratios of Abf2 concentration, *a*, and *n*, result in a stabilization of mtDNA, asymptotically approaching complete stability. **d)** Hemizygous deletions of *MIP1* or *ABF2* in diploid strains are modeled by reducing the concentrations *m* or *a*, respectively, to 50%. Depending on the model parameters, single hemizygotes have mtDNA copy numbers between 50% to 100% of the wild-type. Independent of the model parameters, a double hemizygote always has a mtDNA copy number reduced to 50%.

Similarly, we model mtDNA degradation as a process inhibited by increasing concentrations of Abf2, *a*, such that 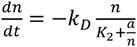, where *k_D_* and *K_2_* are again constants (Fig. 5a,c). Accounting for cell growth by assuming dilution of mtDNA according to exponential growth with a doubling time *T*, we can then balance replication, degradation, and dilution to obtain 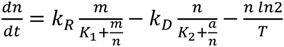 (Fig. 5a). Assuming steady-state, 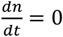, we then obtained an equation directly linking the concentration of mtDNA to that of Mip1 and Abf2.

### Nuclear encoded mtDNA maintenance machinery can couple mtDNA amount to cell volume

One direct implication of this result is that if the amounts of Mip1 and Abf2 increase in direct proportion to cell volume, thereby maintaining constant concentrations *a* and *m*, the steady state solution for *n*, *i.e.* the concentration of mtDNA, is also independent of cell volume. In other words, the mtDNA copy number increases in direct proportion to cell volume (Fig. 5a). Thus, our simple model explains how an increasing amount of mtDNA maintenance machinery in bigger cells can couple mtDNA copy number to cell volume.

Next, we asked whether our model also explains the reduction of mtDNA observed in diploids hemizygous for *MIP1* or *ABF2*. In the model, deletion of one of the alleles of *MIP1* and *ABF2* can be accounted for by reducing *m* or *a*, respectively, by a factor of 2. Solving the steady state model, we then found that the effect of the hemizygous deletions strongly depends on the exact parameters chosen: We can find parameters, for which both hemizygotes cause a reduction to about 70% of mtDNA, reflecting our experimental results (Fig. 5d). However, faster replication (increased *k_R_*) will shift the system to a regime where *MIP1* hemizygotes show almost a 50% reduction, and *ABF2* hemizygotes have nearly unchanged mtDNA amounts. In contrast, increasing the degradation rate *k_D_* results in the opposite behavior. Thus, while our model can be consistent with our experimental findings, single hemizygote deletions are not well-suited to test the model. However, we noticed that our model makes a prediction for a simultaneous manipulation of Mip1 and Abf2 concentrations: Independent of the parameters chosen, if *m* and *a* are changed by the same factor, also the concentration of mtDNA, *n*, follows proportionally (Fig. 5d).

To directly test this prediction, we constructed a strain hemizygous for both *MIP1* and *ABF2*. We verified with qPCR that in this double hemizygote, expression of *MIP1* and *ABF2* is reduced to 50% (Supplementary Fig. 7b-d). As predicted by the model, we find that in this strain also the concentration of mtDNA is reduced close to 50% (Fig. 6a). Since the model also predicts that the effect of the hemizygous deletions should be independent of cell volume, we then repeated the experiments with a Whi5-inducible strain. Consistent with the model, we found that both single- and double hemizygotes show mtDNA copy numbers that increase with cell volume, with similar relative reduction of mtDNA by the hemizygous deletions at all cell volumes (Fig. 6b).

**Figure 6.**
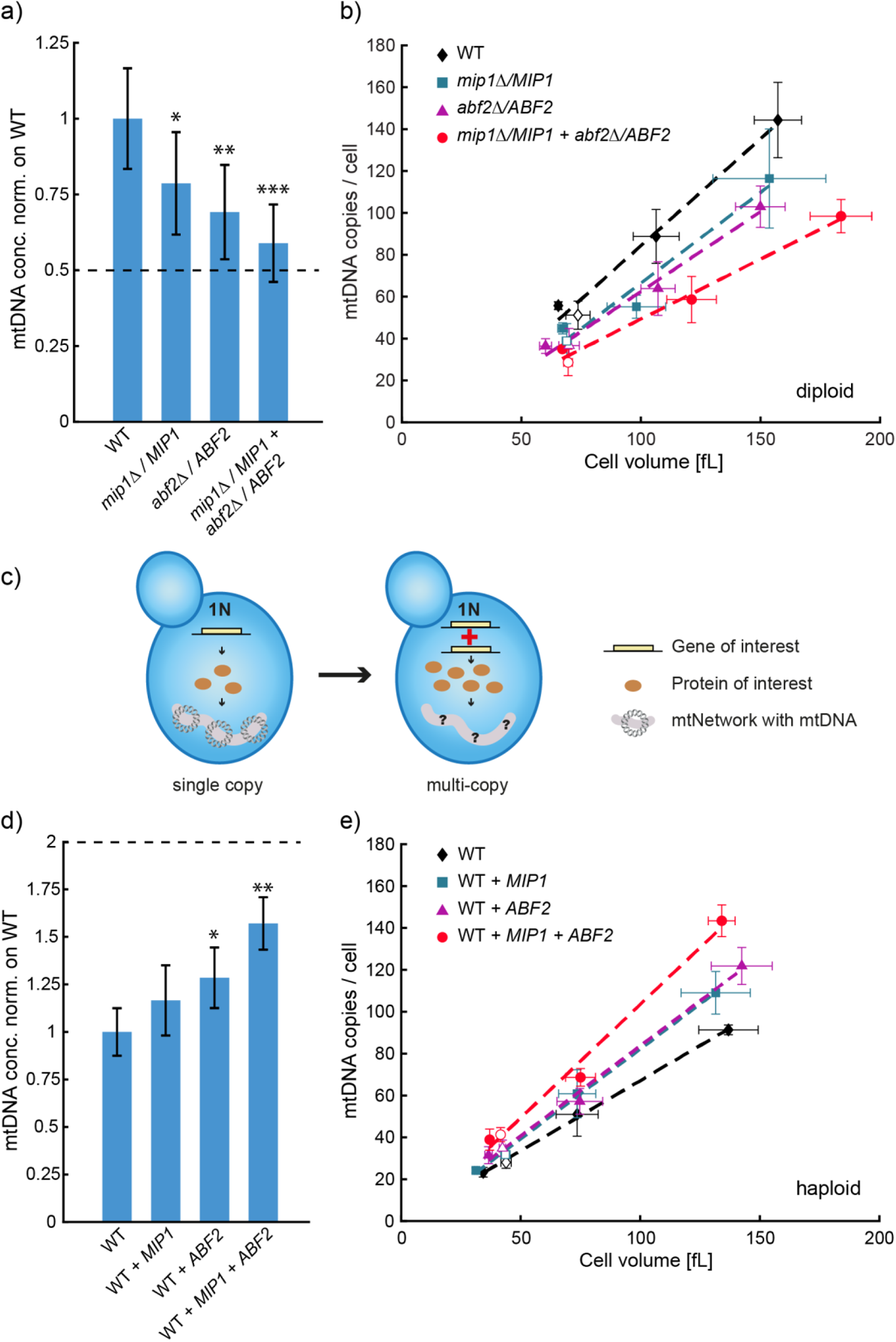
mtDNA copy number is modulated by concentrations of mtDNA maintenance factors. **a)** mtDNA copy numbers per cell as determined by DNA-qPCR of single (data from Fig. 4c) and double hemizygous *MIP1* and *ABF2* strains normalized on wild-type. **b)** mtDNA copy number per cell determined by DNA-qPCR in non-inducible (data from b) and Whi5-inducible hemizygous strains. Error bars indicate standard deviations. Lines indicate linear fits of the means. **c-e)** Additional copies of *MIP1* and/or *ABF2* were endogenously integrated into wild-type and Whi5-inducible haploid strains. mtDNA copy number was determined by DNA-qPCR and mean cell volume was measured with a Coulter counter. mtDNA copy number per cell normalized on wild-type are shown for the non-inducible strains in **d**. mtDNA copy numbers per cell for Whi5-inducible and non-inducible strains are shown in **e**. Significances were determined by a two-tailed t-test (* p<0.05, ** p<0.01,*** p<0.001).

Similar to a simultaneous reduction of Mip1 and Abf2 concentrations, our model also predicts that the mtDNA concentration should increase 2-fold if both Mip1 and Abf2 are 2-fold overexpressed. To test this hypothesis, we constructed haploid strains in which we endogenously integrated additional copies of the *MIP1* and/or *ABF2* genes (Fig. 6c), resulting in a 2-fold increase of *MIP1* and *ABF2* expression, respectively (Supplementary Fig. 8). Consistent with the effect of hemizygous deletions, we find that overexpression of either Mip1 or Abf2 results in a moderate increase of mtDNA concentration, and simultaneous overexpression of both has an additive effect (Fig. 6d). Again, repeating the experiment in a Whi5-inducible strain revealed that the proportional scaling of mtDNA amount with cell volume is maintained in each strain (Fig. 6e). Importantly, however, simultaneous 2-fold overexpression of Mip1 and Abf2 only results in a 57% increase of mtDNA, which is less than the 2-fold increase predicted by our simple model. This suggests that upon overexpression of the most limiting factors for mtDNA maintenance (Mip1 and Abf2), other proteins, which are not included in our model, become limiting. Given our analysis of the expression of mtDNA maintenance factors (Fig. 3), it seems likely that also the amount of those additional factors increases in proportion to cell volume. In contrast to the selective overexpression of only Abf2 and Mip1 we achieved through the additional gene copies, a 2-fold increase of cell volume would then still maintain Mip1 and Abf2 as major limiting factors, coupling mtDNA copy number to cell volume.

## Discussion

In summary, we find that in budding yeast, mtDNA copy number is tightly coupled to cell volume, both in arrested cells growing in G1 and in asynchronously cycling cell populations. This is consistent with early work showing that mtDNA amount per cell increases during a G1 arrest (Petes and Fangman, 1973; Conrad and Newlon, 1982) and with the cell volume of stationary cells (Lee and Johnson, 1977). Because the coupling of mtDNA copy number to cell volume can maintain constant mtDNA concentrations independent of cell cycle stage, it provides an elegant mechanism for cells to maintain mtDNA homeostasis during cell growth, without requiring a coordination of mtDNA replication with the cell cycle. Interestingly, this strategy for DNA maintenance is opposite to that implemented by eukaryotes to maintain nuclear DNA, where DNA replication is strictly coupled to cell cycle progression but not directly linked to cell volume.

Based on our experiments and theoretical considerations, we propose a molecular mechanism that can quantitatively explain the increase of mtDNA amount with cell volume: mtDNA concentration is determined by the rates of replication, degradation and dilution by cell growth. Both replication and mtDNA stability can depend on mtDNA maintenance factors in a dose-dependent manner. Larger cells have overall more proteins, and also have a higher amount of mtDNA replication and maintenance factors. As a consequence, larger cells can maintain a higher mtDNA copy number (Fig. 7).

**Figure 7.**
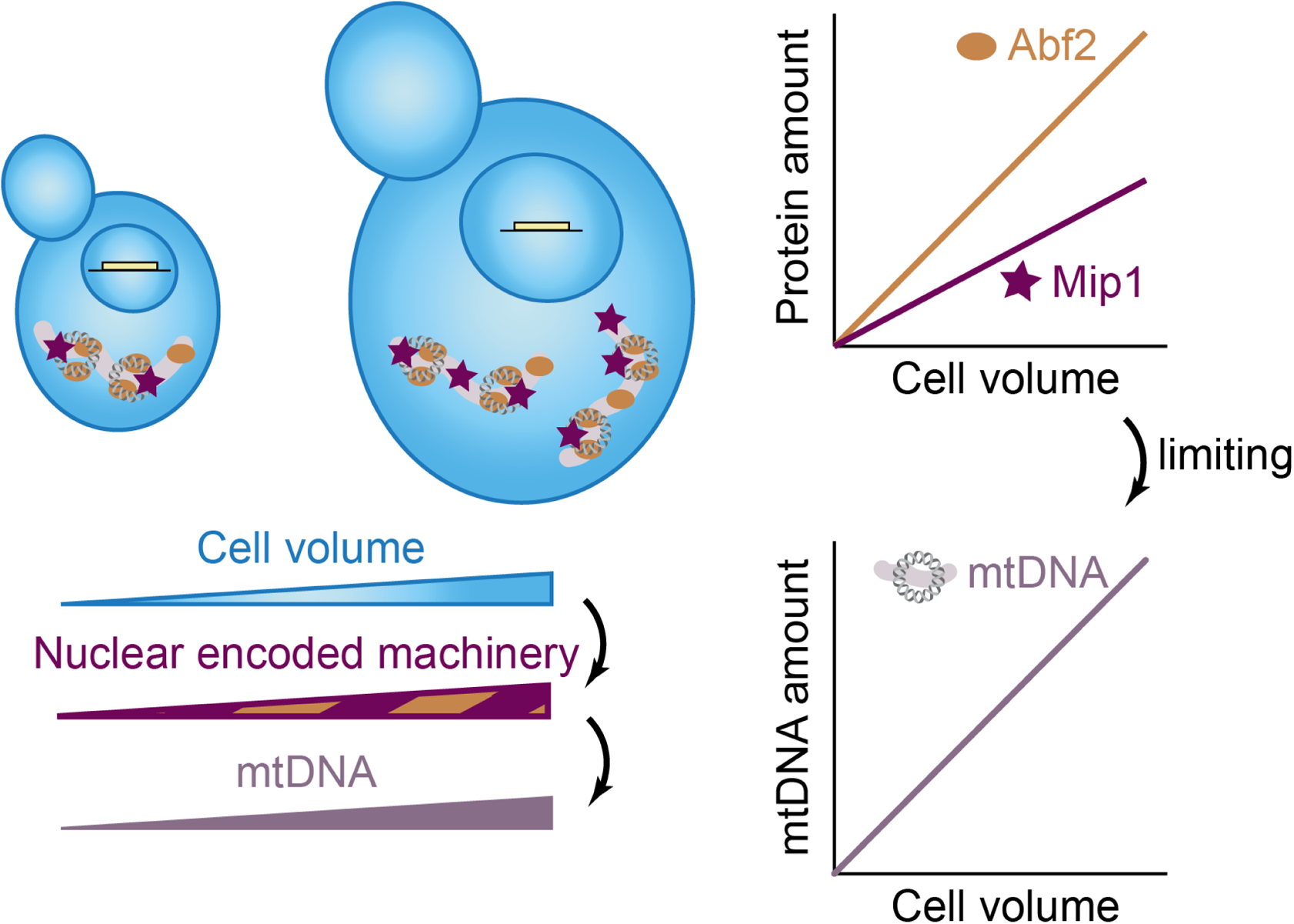
Illustration of the limiting-machinery mechanism for mtDNA homeostasis during cell growth. Cell volume sets the abundance of the nuclear encoded mtDNA maintenance machinery by global regulation of gene expression. This includes the mitochondrial DNA polymerase Mip1 and the packaging factor Abf2, whose amount increases with cell volume, and whose abundance has a direct impact on mtDNA amounts. Thus, controlling limiting mtDNA maintenance factors by global regulation of gene expression with cell volume provides a mechanism of how mtDNA homeostasis is achieved during cell growth.

A single perfectly limiting component of the mtDNA maintenance machinery could directly lead to mtDNA amounts increasing in proportion to cell volume; for example, if larger cells had proportionally more mtDNA polymerase, and mtDNA replication rate increased in proportion to polymerase number.

However, our experiments suggest that no single protein is perfectly limiting (such that a doubling of protein concentration would lead to a doubling of mtDNA copy number). Instead, we identified two partially limiting factors, Mip1 and Abf2. To obtain a conceptual understanding of how several partially limiting factors, each of which is nuclear encoded and increasing in amount with cell volume, affect mtDNA numbers, we built a minimal mathematical model describing the contribution of mtDNA replication, degradation and dilution by growth. In the model, replication and degradation are promoted and prevented, respectively, by the most limiting factors Mip1 and Abf2. We find that even in a situation where neither Mip1 nor Abf2 are perfectly limiting, they synergistically cause an increase in mtDNA amount proportional to cell volume, thus maintaining constant concentrations. The model correctly predicts the ∼50% decrease of mtDNA amount observed in diploids hemizygous for *MIP1* and *ABF2*. This suggests that in the situation where the most limiting factors Mip1 and Abf2 are both reduced below wild-type levels, all other factors whose concentrations are not reduced are present in excess and thus have only a weak additional limiting contribution to mtDNA maintenance. By contrast, upon 2-fold overexpression of Mip1 and Abf2, we observe a stronger deviation from the simple model, which is consistent with other factors becoming partially limiting. At this point, we do not think it is helpful to extend our simple model to many additional factors because the exact molecular mechanisms of their contribution, including rate constants, are not known. However, it seems likely that adding also additional gene copies of the other limiting factors will result in a further increase of mtDNA concentration, eventually reaching a 2-fold increase. Since increasing cell volume likely causes an increasing abundance of all limiting factors, their combined effect is needed to fully explain the coordination of mtDNA amount and cell volume.

Importantly, while our work identifies a mechanism that couples mtDNA amount to cell volume, additional mechanisms can modulate mtDNA homeostasis. For example, additional mechanisms may be responsible to achieve adaptation to changing environments, such as the reduction of mtDNA copy number we observed for cells grown on fermentable compared to non-fermentable media. Within the framework of the model, this can be achieved by regulating the concentration of limiting factors, but also by changing any of the rate constants. For example, Mrx6 was proposed to modulate mtDNA concentration through its role in degradation of factors involved in mtDNA replication (Göke et al., 2020). Moreover, while our proposed mechanism achieves mtDNA homeostasis without cell-cycle-dependent regulation, and we observe the increase of mtDNA amount with cell volume both in G1-arrested and in asynchronous cell populations, we cannot exclude modest cell-cycle-modulation. In this context it is noteworthy that in parasitic kinetoplastids such as *T. brucei* (Woodward and Gull, 1990), mtDNA replication is strongly coupled to the cell cycle, suggesting that also other organisms could employ cell cycle regulation as an additional level of regulation. In fact, in budding yeast both *MIP1* and *ABF2* expression exhibit weak cell-cycle-dependence (Santos et al., 2015), which would propagate to mtDNA amounts in our model.

Our study reveals that in addition to mitochondrial network volume (Rafelski et al., 2012), also mtDNA amount increases with cell volume. In addition, we did not observe major changes of mitochondrial diameter and ultrastructure with increasing cell volume, which is consistent with a previous report that observed only a weak dependence of mitochondrial structure on cell volume in mouse cells (Miettinen et al., 2014). However, even in the absence of major structural changes, mitochondrial function, including respiratory activity, might be modulated by cell size. For example, previous work in mammalian cells found that while mitochondrial mass increases with cell volume, mitochondrial function is optimal at intermediate cell volumes (Miettinen et al., 2014; Miettinen and Björklund, 2016). Similarly, molecular reorganization could modulate mitochondrial function in yeast such that optimal function is achieved at intermediate volumes.

In essence, the mechanism we propose here for mtDNA homeostasis in budding yeast only requires that the limiting components of the mtDNA maintenance machinery increase in abundance with cell volume. One key feature of this mechanism is that it is robust towards perturbations and fluctuations in mtDNA concentration. Since steady state concentrations of mtDNA are set by the concentrations of the nuclear encoded limiting factors, which are themselves independent of mtDNA concentration, cells with excess or low levels of mtDNA will regress back to the stable steady state without a need for an active feedback mechanism. Indeed, such passive regression to the mean has been observed for the nucleoid number in fission yeast (Jajoo et al., 2016).

We anticipate that mtDNA homeostasis achieved through limiting nuclear encoded machinery is a regulatory principle that is conserved across eukaryotes. However, the identity of the most limiting factors might vary between organisms. For example, it has been shown that in animals, the Abf2 homolog TFAM (Larsson et al., 1998; Matsushima et al., 2003; Ekstrand et al., 2004) as well as the mitochondrial helicase Twinkle (Tyynismaa et al., 2004) have a strong dose-dependent effect on mtDNA copy number. Further supporting our hypothesis, a recent study revealed that many mitochondrial proteins, including TFAM, increase in amount with increasing volume of human epithelial cells (Lanz et al., 2021). More generally, the increase of global protein amounts with increasing cell size due to increased biosynthetic capacity is widely conserved across eukaryotes. Thus, limiting nuclear encoded genes provide a robust mechanism to achieve mtDNA homeostasis in growing cells.

## Methods

### Yeast strains

All yeast strains used in this work are derived from W303 and listed in Supplementary Table 2. Construction of yeast strains was performed with standard methods. Transformants were verified by control PCRs and sequencing.

Microscopy strains (haploid, diploid, ρ^0^) were generated from parental strains containing LacO-arrays in mtDNA (yCO380, yCO381) (Osman et al., 2015). Endogenous *WHI5* was deleted in yCO380 and β-estradiol-inducible *WHI5* was integrated (KSE113-1) followed by endogenous integration of a plasmid carrying the β-estradiol transcription factor (FRP880) (Ottoz et al., 2014), resulting in strain ASY11-2B.

Next, the plasmid ASE001-5 containing *mKate2* and *LacI* tagged two copies of *mNeon* was integrated into the HO-locus to obtain strain ASY13-1. To generate the diploid strain ASY15-1, yCO381 was transformed with ASE001-5 and crossed with ASY11-2B.

### Yeast culturing

All strains were grown at 30°C in a shaking incubator at 250 rpm (Infors, Ecotron).

Prior to growing cells on non-fermentable medium (synthetic complete media containing 2% glycerol and 1% ethanol, SCGE), strains were grown for at least 6 h on YPD. Then, cells were washed with SCGE and transferred into SCGE. Cultures were grown for about 24 h in exponential phase when directly used for experiments, or grown for at least 12 h before adding β-estradiol to Whi5-inducible strains. To tune Whi5 concentration in Whi5-inducible strains, cells were grown for another 24 h in the presence of the respective β-estradiol concentration. For haploid strains, concentrations of 0 nM, 10 nM, and 30 nM, and for diploids 0 nM, 15 nM and 60 nM β-estradiol were used.

When using fermentable medium, cells were directly inoculated in synthetic complete media containing 2% dextrose (SCD) and grown for at least 12 h prior addition of β-estradiol. For both haploid and diploid strains, concentrations of 0 nM, 15 nM, 60 nM and 150 nM β-estradiol were added, and cells grown for additional 24 h.

For the G1-arrest (Fig. 1c), a *cln1/2/3* deletion strain with Cln1 expressed from a β-estradiol-inducible promoter was used (Ewald et al., 2016). Prior G1-arrest, cells were grown at least 6 h on YPD with 30 nM β-estradiol, transferred into SCGE with 30 nM β-estradiol and grown for about 24 h. For fermentable conditions, cells were directly inoculated in SCD medium with 60 nM β-estradiol and grown for 24 h. To initiate the G1 arrest, cells were washed with the respective medium without hormone, and cultures were then harvested every hour (SCGE 0-8 h; SCD 0-6 h).

Steady-state exponential growth conditions were obtained by regularly measuring optical densities using a spectrophotometer (NanoDrop One^C^, Thermo Fisher Scientific) and ensuring OD_600_ < 1 through appropriate dilutions. To determine mean cell volumes of cell populations, cell volume distributions were measured using a Coulter counter (Beckman Coulter, Z2 Particle Counter) after sonication. Due to technical limitations of the measurable ranges, samples were measure twice with two different settings (Range 1: 10 fL - 328 fL, gain: 256, current: 0.707 ma; Range 2: 328 fL – 1856 fL, gain: 256, current: 0.125 ma). We then used both measurements to calculate a mean volume within the combined cell volume range.

### mtDNA copy number measurements

Cells were cultivated in 50 ml of the respective medium with corresponding β-estradiol concentrations. Prior to harvesting, cell volume distributions and optical density were measured. Cell cultures were spun at 4000 rpm and pellets were washed with 1 ml double-distilled water.

gDNA was extracted by phenol-chloroform-isoamyl alcohol (PCI) extraction. More precisely, cells were mechanically disrupted by vortexing at 3000 oscillations per minute (Mini-BeadBeater 24, 230V, BioSpec Products) with glass beads in 200 µl DNA extraction buffer pH 8.0 (2% TritonX100, 1% SDS, 100 mM NaCl, 10 mM TRIS, 1 mM EDTA) and 200 µl PCI. After centrifugation at 13,000 rpm, the aqueous phase was taken to precipitate gDNA with 500 µl 100% EtOH, centrifugation was repeated and the pellet was then washed with 800 µl 70% EtOH.

To remove RNA residues, the pellet was solved in nuclease-free water, treated with 1 mg/ml RNase A (DNase-free) and incubated for 30 min at 37°C. Subsequently, DNA extraction buffer and PCI were added and extraction steps were repeated. DNA concentrations were determined with a spectrophotometer (NanoDrop One^C^, Thermo Fisher Scientific) through measurements at 260 nm. For qPCRs, 1 ng DNA was used.

Next, quantitative PCR was performed on a LightCycler 480 Multiwell Plate 96 (Roche). For amplification, a DNA-binding fluorescent dye (BioRad, SsoAdvanced Universal SYBR Green Supermix) and specific primers for nuclear DNA (nDNA) genes *ACT1*, *MIP1* and *MRX6* and mtDNA genes *COX2* and *COX3* (Supplementary Table 3) were used. For strains in which *MIP1* gene copy was manipulated, *MIP1* primers were omitted from the analysis. Note that the initial denaturation time was set to 10 min. Each sample was measured in technical triplicates. For further analysis, mean Cq-values of the technical replicates were used. Single technical replicates were excluded from the analysis when the standard deviation was higher than 0.5.

To correct for differences in primer efficiencies and enable absolute measurements of DNA concentrations, a calibration standard was obtained by constructing a single PCR product containing all amplified sequences. A standard dilution series with defined input concentrations (1 pg/µl - 10^-4^ pg/µl) was then performed to obtain a standard curve for each primer pair. A linear fit to these calibration curves was finally used to calculate concentrations from qPCR measurements (Supplementary Fig. 9).

Concentrations of each gene were calculated, and nDNA concentrations (based on *ACT1*, *MIP1*, *MRX6*) and mtDNA concentrations (*COX2*, *COX3*) were pooled by calculating the mean, respectively. mtDNA concentrations were then normalized on nDNA, to obtain the relative mtDNA copy number per nDNA. By performing bud counts, a budding index (percentage of budded cells) was determined for each cell population, and used to calculate the average nDNA amount per cell: 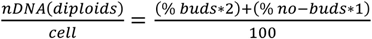 or 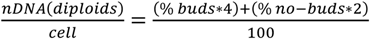. Multiplication with mtDNA copies per nDNA then allowed us to determine the average mtDNA copy number per cell.

### mRNA measurements

RNA samples in Fig. 3a were taken from experiments performed in the study of Claude et al. (Claude et al., 2021), Fig. 2 b-c. Briefly, cells were cultivated in 25 ml of the respective media and grown as described above. RNA was extracted by a hot acidic phenol (Sigma-Aldrich) and chloroform (Thermo Fisher Scientific) extraction. RNA extractions in Supplementary Fig. 1, 5, 7, 8 were performed with the YeaStar RNA Kit (Zymo Research) following the instructions of the given protocol. DNA contaminations were removed by a DNA digestion step using DNaseI (Life Technologies). cDNA was synthesized by using 1000 ng total RNA, random primers and following the protocol of the high-capacity cDNA reverse-transcription kit (Thermo Fisher Scientific). mRNA expression levels of *MIP1*, *ABF2*, *PIM1*, *MTF1*, *RPO41* and *MRX6* were measured by qPCR using the fluorescent dye SybrGreen for detection. 2 µl of a 1:10 dilution of cDNA was used, except for the ribosomal RNA *RDN18*, for which 2 µl of a 1:200 dilution was used. Each sample was measured in triplicates and concentrations were calculated after normalization on *RDN18*.

### Analysis of transcript and protein cell-size-dependence based on Swaffer et al

To compare the cell-size-dependence of the transcripts of mtDNA maintenance factors (*ABF2*, *HMI1*, *MGM101*, *MHR1*, *MIP1*, *MRX6*, *MTF1*, *PIF1*, *PIM1*, *RAD53*, *RIM1*, *RPO41*, *RRM3*) with that of scaling control genes (*ACT1* and the RNA polymerase II subunits *RPB2*, *RPB3*, *RPB4*, *RPB5*, *RPB7*, *RPB8*, *RPB9*, *RPB10*, *RPB11*, *RPO21*) and the sub-scaling histones (*HHF1*, *HHF2*, *HHO1*, *HHT1*, *HTA1*, *HTA2*, *HTB1*, *HTB2*, *HTZ1*), we analyzed two data sets published by Swaffer et al. (Swaffer et al., 2021a). For the first data set, budded cells were sorted into four different size bins using a total protein stain as a measure for cell size and then analyzed with RNA sequencing. We compared the ratio (mean of two independent replicates) between the relative expression levels in the largest and smallest cells. For the second data set, cells were elutriated and arrested in G1 for different amounts of time before synchronous release, resulting in different cell volumes at the time of cell cycle entry. The temporal evolution of the transcriptome during cell cycle progression was then analyzed with RNA sequencing. The relative expression throughout the cell cycle was then calculated as the Area Under the Curve (AUC) of the expression level time course (after applying a spline). Again, we compared the ratio between the largest and smallest cells (mean of two independent replicates). Based on the combined analysis of both datasets, all mtDNA factors we analyzed were classified as ‘scaling’ by Swaffer et al. In the first data set (but not the second) *RAD53* showed a strongly increased expression in big cells, which we attributed to its strong cell-cycle-dependence. We therefore excluded *RAD53* from further analysis.

Of the mtDNA maintenance factors described above, Abf2, Mhr1, Pim1 and Mhr1 are included in the analysis of the dependence of protein amount on cell volume by Swaffer et al. based on flow-cytometry measurement on strains carrying GFP-tagged alleles of the respective proteins performed by Parts et al. (Parts et al., 2014). Briefly, the normalized slope of GFP intensity as a function side scatter was used to estimate the cell-volume-dependence of the protein amount. Proteins maintained at a perfectly constant amount would be expected to exhibit a slope of 0, while proteins maintained at a constant concentration would exhibit a slope of 1. Parts et al. performed two independent biological replicates, which we analyzed separately.

### Microscopy

For imaging, coverslips (µ-Slide 8 Well, ibi-Treat, ibidi) were covered with 200 µl Concanavalin A (ConA, 1mg/ml in H_2_O) and incubated for 5-10 min. The wells were then washed twice with water and left to air dry.

Cells were cultivated as described in 5 ml medium. 1 ml of the culture was spun at 13.000 rpm and washed twice with 1 ml medium. Depending on the OD, the pellet was dissolved in 200 µl - 500 µl of the respective medium and 200 µl were transferred to the ConA covered well. The cells were then allowed to settle down for about 5 min, before the supernatant was removed and the wells were washed twice with medium. 200 µl medium were used to cover the wells.

Live-cell fluorescence microscopy experiments were performed on a Zeiss LSM 800 microscope (software: Zen 2.3, blue edition) equipped with a scanning disk confocal and an axiocam 506 camera, using the confocal mode. Images were taken using a 63x /1.4 Oil DIC objective. Z-stacks were acquired over 15.05 µm in 0.35 µm increments. mKate2 was imaged with an excitation wavelength of 561 nm, and detecting emission between 610-700 nm. mNeon was excited at 488 nm and detected between 410-546 nm. Bright field images were taken using the transmitted light detector (T-PMT).

### Cell segmentation

Cell segmentation was performed using Cellpose v0.6 (Stringer et al., 2021) with the ‘cell diameter’ parameter set to 50 pixels, ‘flow threshold’ set to 0.4, and ‘cell probability threshold’ set to 0. Cell-ACDC (Padovani et al., 2021) was used to manually correct segmentation, annotate buds to their corresponding mother cells, and calculate cell volume.

### Nucleoids counting and mitochondrial network volume calculation

To count the number of nucleoids and compute the mitochondrial network volume from confocal 3D z-stack images, we developed a custom routine written in Python.

The analysis steps are the following: 1) Application of a 3D gaussian filter with a small sigma (0.75 voxel) of both the nucleoids and mitochondria signals; 2) instance segmentation of the mitochondria signal using automatic Li thresholding (Li and Tam, 1998) (*threshold_li* function from the library *scikit-image* (van der Walt et al., 2014)); 3) normalization of the mitochondria signal using the median of the voxel intensities classified as mitochondria in step 2; 4) 3D local maxima detection (peaks) in the nucleoids signal using the *peak_local_max* function from the library *scikit-image*; 5) discarding of peaks that are below a threshold value determined with the automatic Li thresholding algorithm; 6) discarding of overlapping peaks: if two or more peaks are within a resolution limited volume only the peak with highest intensity is retained. The resolution limited volume is determined as a spheroid with *x* and *y* radii equal to the Abbe diffraction limit and *z* radius equal to 1 µm. With a numerical aperture of 1.4 and mNeon emission wavelength of about 509 nm, the resolution limited volume has *x* = *y* = 0.222 µm radius; 7) the remaining peaks undergo a subsequent iterative filtering routine: a) each voxel classified as mitochondria in step 2 is further classified as inside or outside of the nucleoids. A voxel is outside of the nucleoid if it is not within the resolution limited volume centred at the peak coordinates; b) the nucleoids signal is normalized by the mean of the voxel intensities classified as outside of the nucleoids (step 7a); c) the normalized intensity distribution of the voxels inside each nucleoid volume is compared to the same voxels from the mitochondria signal. The comparison is performed with a Welch’s t-test and if the *p-*value is above 0.025 or the t-statistic is negative (*i.e.*, mitochondria signal higher then nucleoid’s signal) the peak is discarded; d) steps a) to c) are repeated until the number of nucleoids stops changing. The assumption of comparing the nucleoids signal to the mitochondria signal is that a peak is a valid nucleoid only if it has an intensity significantly higher than the corresponding mitochondria signal (after normalization).

The resulting peaks are considered valid nucleoids and are therefore counted. The mitochondrial network volume is computed as the sum of the voxels classified as mitochondria in step 2. Note that due to the optical resolution limit, the width of the network is not measured accurately with confocal microscopy and the obtained mitochondria network volume is therefore not an absolute measure for the physical volume of the mitochondria.

### Electron microscopy

For cell volume quantification, a sample was taken from each culture before the cells were prepared for electron microscopy. These samples were analyzed by light microscopy and DIC images of living cells were taken using a Zeiss Axiophot microscope equipped with a Plan-Neofluar 100x/1.30 Oil objective (Carl Zeiss Lichtmikroskopie, Göttingen, Germany) and a Leica DFC360 FX camera operated with the Leica LAS AF software version 2.2.1 (Leica Microsystems, Wetzlar, Germany). Cell segmentation and volume estimation were performed using Cell-ACDC (Padovani et al., 2021) as described above.

Fixation of yeast cells for electron microscopy with glutaraldehyde and potassium permanganate was performed as described in (Perkins and McCaffery, 2007) with the following changes: Glutaraldehyde fixation was performed with 3% glutaraldehyde, 0.1 M sodium cacodylate, 1 mM CaCl_2_, pH 7.2 and the samples were subsequently washed with 0.1 M sodium cacodylate, 1 mM CaCl_2_, pH 7.2. Treatment with potassium permanganate was either performed before (cells grown in SCD) or after (cells grown in SCGE) embedding in agar. After treatment with sodium metaperiodate and overnight staining with 2% uranyl acetate at room temperature, dehydration of chemically fixed yeast cells with ethanol and propylene oxide, Epon infiltration, and contrast enhancement of ultrathin sections were essentially performed as described in (Unger et al., 2017) with the following modifications: All dehydration steps were performed at 4 °C, Epon infiltration was performed at room temperature, and contrast enhancement of ultrathin sections was performed for 15 min with 2% uranyl acetate and for 3 min with lead citrate. Electron micrographs were taken using a JEOL JEM-1400 Plus transmission electron microscope operated at 80 kV, a 3296×2472 pixels JEOL Ruby CCD camera, and the TEM Center software, either Ver.1.7.12.1984 or Ver.1.7.19.2439 (JEOL, Tokyo, Japan). As an estimate of mitochondrial diameter the length of the minor axis of 100 mitochondria was measured for each sample from electron micrographs using Fiji (Schindelin et al., 2012).

### Flow cytometry

Wild-type cells in which Mip1 and Abf2, respectively, were endogenously tagged with mCitrine were analyzed with flow cytometry to determine the dependence of Mip1 and Abf2 protein amounts on cell volume.

Cells were cultured as described above. After 16-20 h of growth on SCGE, cultures were diluted, split into 3 technical replicates and for control measurements shown in Supplementary Fig. 6a-b, ß-estradiol was added. Optical density was measured with a spectrophotometer (NanoDrop One^C^, Thermo Fisher Scientific) and only cultures with an OD_600_ <0.9 were included in the flow cytometry measurements. Cultures were kept on ice until measurement. After sonification for 10 sec mean cell volume of each culture was determined using a Coulter counter. The Flow Cytometry measurement was performed on a CytoFlex S Flow Cytometer (Beckman Coulter) and the parameters FSC-A, SSC-A and total fluorescence intensity using the FITC channel (excitation at 488 nm and detection with a 525/40 nm filter) were recorded. Cells were analyzed at a slow flow rate (10 µL/min) and data were collected from 50.000 events per sample. Through a standard gating strategy (Supplementary Fig. 6d), cell debris, particles and doublets were excluded from the analysis. Identical settings were used for all measurements. To correct for the autofluorescence of yeast cells, the parent strain without mCitrine tag was measured.

After confirming that differences between technical replicates were negligible, the 3 replicates measured on one day were pooled, and binned according to SSC-A, which is a good proxy for cell volume (Supplementary Fig. 6). To correct for autofluorescence, for each bin the mean signal of the autofluorescence control was subtracted from the mean signal of the fluorescent strain in the same bin. This analysis was repeated for data obtained on a different day (3 technical replicates each). Background-corrected signals obtained on each day were then averaged. For each bin, the maximum (minimum) of the two signals plus (minus) the standard error associated with the measurement of the fluorescent strain was used to obtain an estimate of the experimental error.

### Model

To better understand the effect of limiting mtDNA maintenance machinery on cell-volume-dependent mtDNA concentration, *n*, we built a minimal mathematical model, neglecting cell-to-cell variability and any potential contributions of asymmetric mtDNA inheritance between mother cells and their buds. We assumed that the rate of mtDNA replication is given by the concentrations of mtDNA polymerase Mip1, *m*, as well as the mtDNA concentration such that the synthesis rate can be described by Michaelis Menten like kinetics, 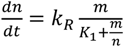. In the limit of saturating Mip1 concentrations, the synthesis rate then approaches the constant *k_R_* multiplied by the concentration of mtDNA. At low Mip1 concentrations, replication is limited by the polymerase Mip1 and thus the synthesis rate increases in direct proportion with *m*. *K_1_* describes the dissociation constant of Mip1 and mtDNA, respectively. In addition, we assume that in the absence of Abf2, each mtDNA molecule is degraded with a rate 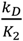, where *k_D_* and *K_2_* are again constants. Increasing concentrations of Abf2, *a*, then stoichiometrically protect mtDNA from degradation, such that the total rate of mtDNA degradation can be described by 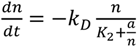.

Finally, we account for the fact that mtDNA is diluted by cell growth by assuming exponential growth with a doubling time *T*. Combining the contributions of replication, degradation, and dilution, we then find that

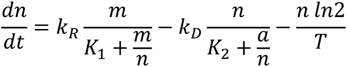

In steady state, we can then assume that 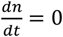, so that

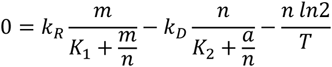

From this equation it can be immediately seen that as long as the concentrations of Mip1 and Abf2 are constant, *i.e.* that the amounts of Mip1 and Abf2 increase in proportion to cell volume, also the concentration of mtDNA is maintained constant, *i.e.* the mtDNA copy number increases in proportion to cell volume.

To understand the impact of hemizygous *MIP1* and *ABF2* deletions, we then chose specific parameters (‘wild-type’: *m* = 5, *a* = 100, *T* = 150, *k_R_* = 0.01 or 0.1, *k_D_* = 1 or 10, *K*_1_ = 5, *K*_2_ = 100), and solved the steady state equation using *Matlab*. Note that while the model is not meant to accurately reflect the quantitative details of budding yeast cells, the parameters are chosen such that the relative ratios of *m*, *a*, and *n* are roughly in the range expected from our measurements and previous estimates (Ghaemmaghami et al., 2003).

## Acknowledgements

We thank Jennifer Ewald for sharing strains, Matthew Swaffer for help with data analysis, Benedikt Westermann and Stefan Geimer for help with electron microscopy, and Max Harner, Johanna Frickel, Simon Schrott, Aylin Göke, and members of the Institute of Functional Epigenetics for discussions. This work was funded by the Deutsche Forschungsgemeinschaft (DFG, German Research Foundation) - 431480687 and 459304237, by the Human Frontier Science Program (career development award to K.M.S.), by the Elitenetzwerk Bayern through the Biological Physics program (T.K.) and the Helmholtz Gesellschaft.

## Notes

### Competing Interest Statement

The authors have declared no competing interest.

### Summary of Updates

The revised manuscript includes EM data on cells grown on media with a non-fermentable carbon source (SCGE).

