## Supplementary Information for "Regulation with cell size ensures mitochondrial DNA homeostasis during cell growth"

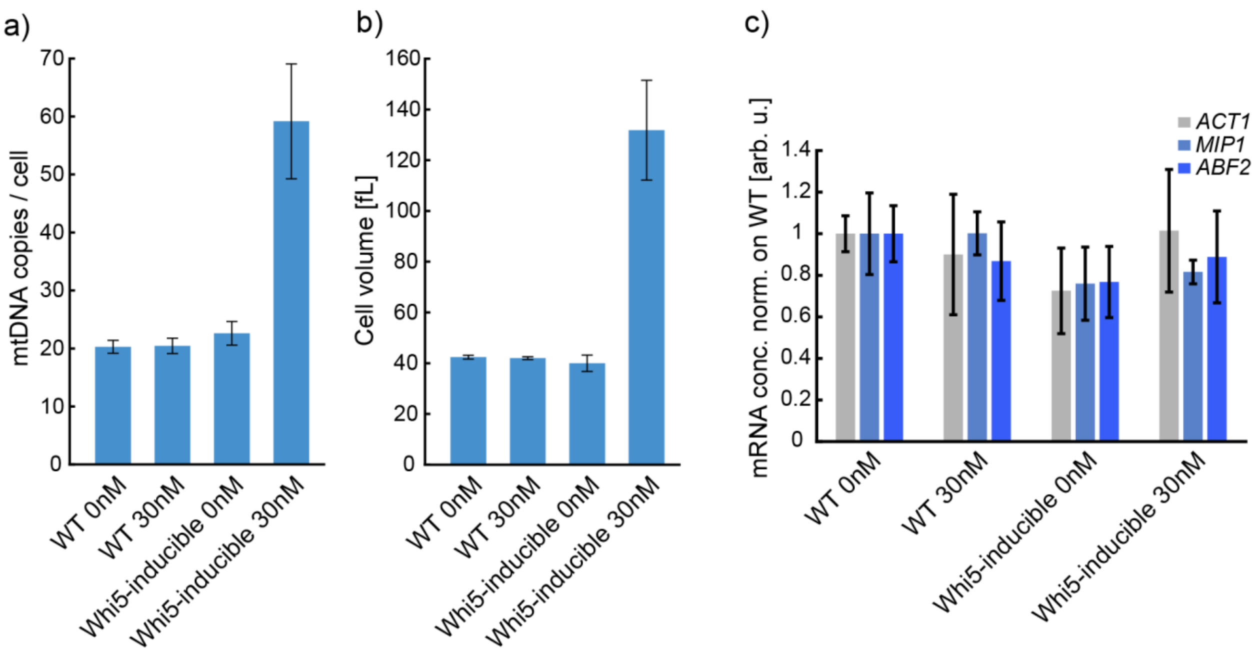


**Supplementary Figure 1: β-estradiol alone does not affect mtDNA copy number (a), cell volume (b) and mRNA levels (c)**. Wild-type (WT) and Whi5-inducible strains were grown on SCGE and treated with β-estradiol as indicated. **a)** mtDNA copy number was determined by DNA-qPCR. No effect of β-estradiol was found for wild-type cells, whereas the Whi5-inducible strain shows a 3-fold increase. **b)** Mean cell volume was measured with a Coulter counter. As for mtDNA, addition of 30 nM β-estradiol causes no changes in cell volume for wild-type cells. **c)** RT-qPCR was performed to measure transcript levels in wild-type. Cq-values were normalized on *RDN18* and mRNA concentrations were normalized on wild-type levels. No significant changes in transcript levels were found after addition of 30 nM β-estradiol for wild-type cells.

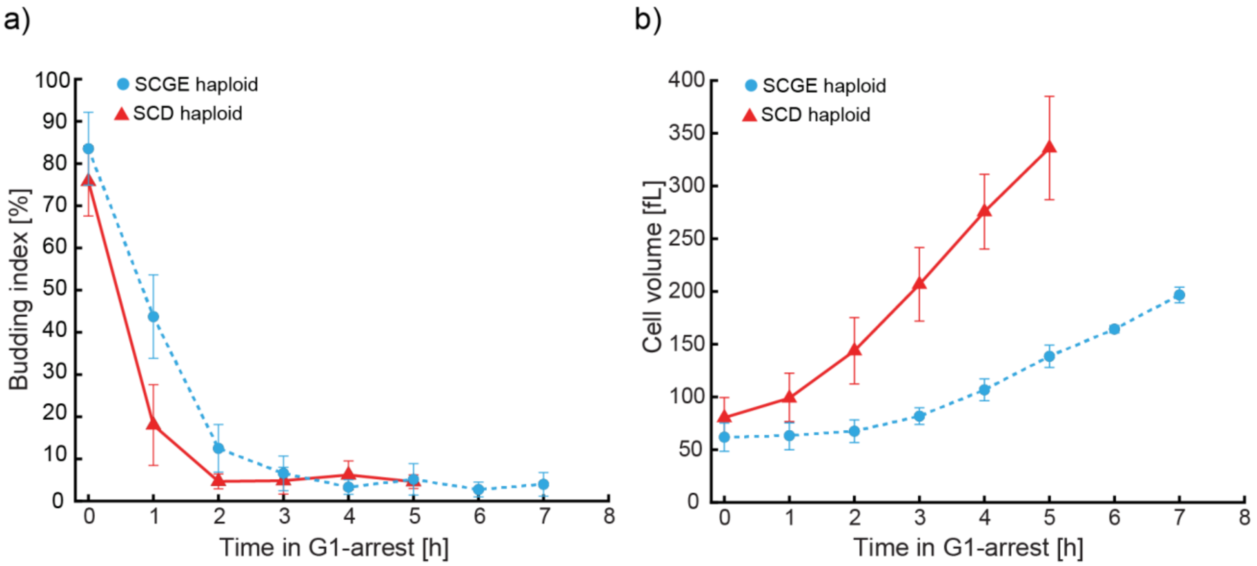


**Supplementary Figure 2: Cell volume and budding index during G1-arrests corresponding to the experiments shown in Fig. 1c. a)** Cell volume was measured with a Coulter counter. **b)** Budding index (fraction of budding cells) was determined by bud counting. After 2 h most cells are arrested in G1, and mean cell volume continuously increases. To account for the slower growth of cells on SCGE, growth was followed for two additional time points.

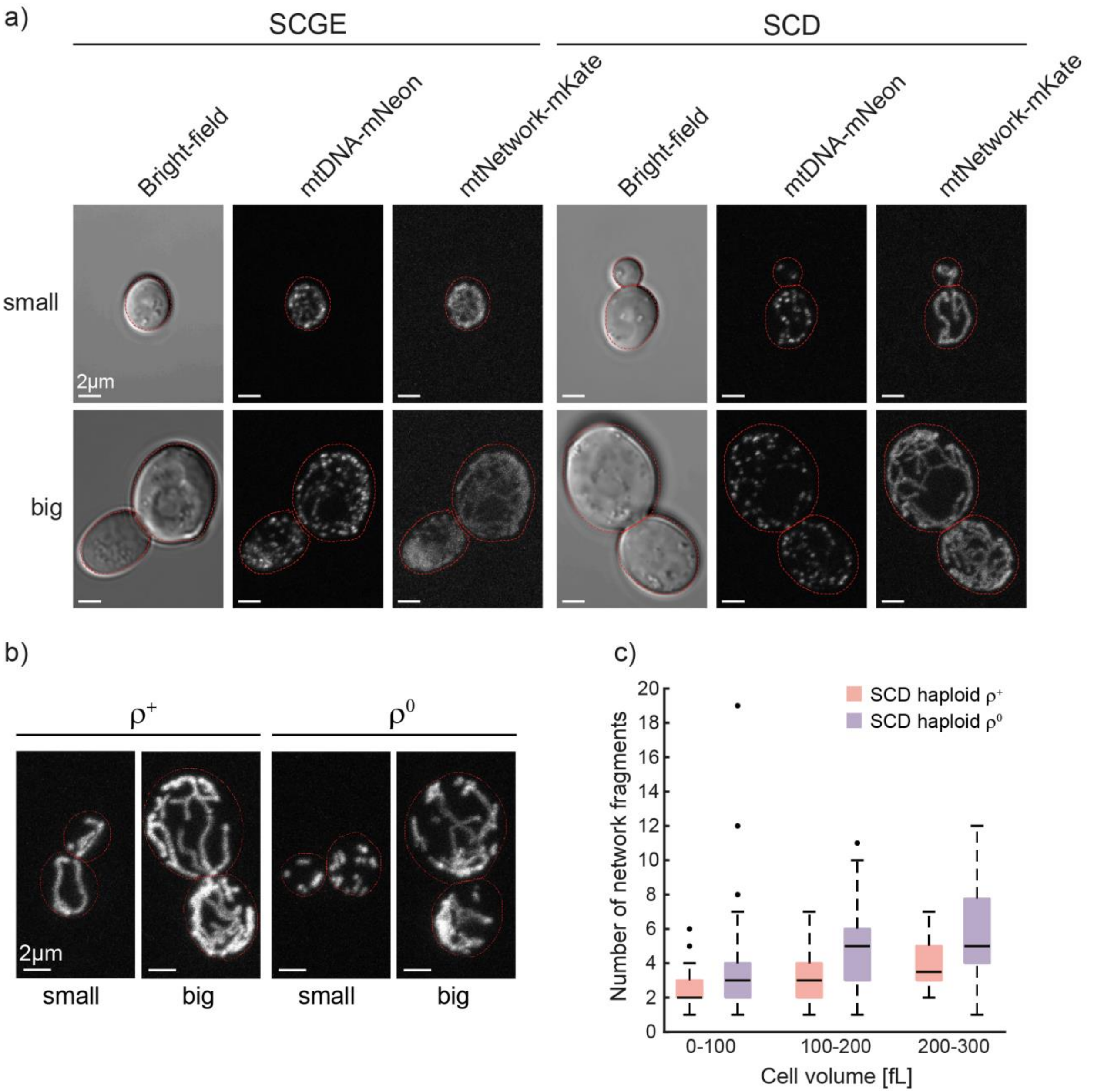


**Supplementary Figure 3: Microscopy images and quantification of network fragments in haploid $\rho^+$ and $\rho^0$ cells.** **a)** Same images as in Fig. 2b are shown without labelling of network segmentation and identified mtDNA. **b)** Same images of haploid $\rho^+$ and $\rho^0$ cells as in Fig. 2e are shown without network segmentation. **c)** The number of separate mitochondrial network fragments in haploid $\rho^+$ and $\rho^0$ cells was calculated for each cell form the mitochondrial network segmentations obtained in Fig. 2e. Cell volume was binned as indicated. Box plots depict median (black line), 25th, 75th percentile (box). Whiskers indicate extreme values still within 1.5 interquartile ranges and outliers are depicted as single black points.

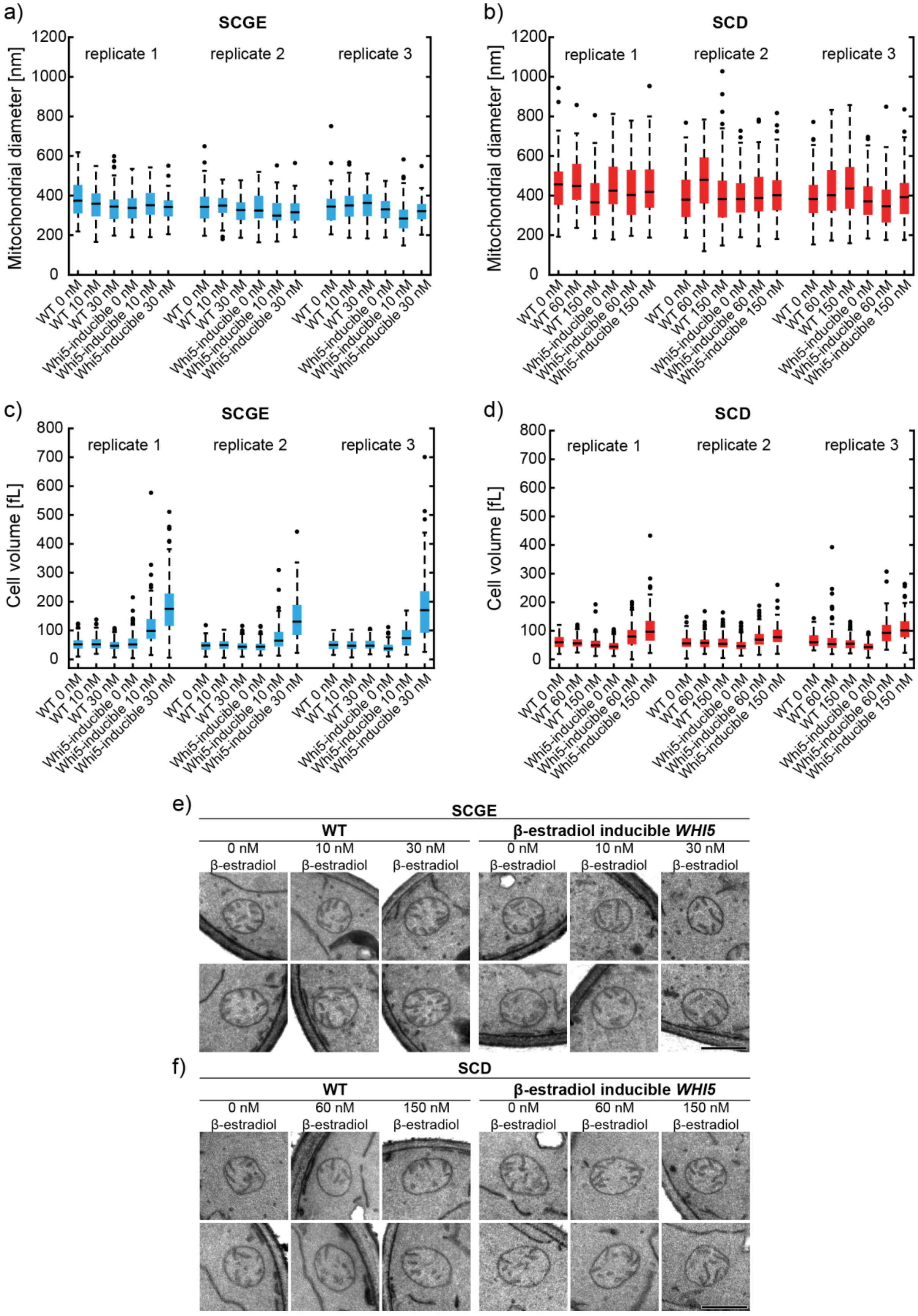
a)
SCGE
Mitochondrial diameter [nm]
0
200
400
600
800
1000
1200
replicate 1
replicate 2
replicate 3
WT 0 nM
WT 10 nM
WT 30 nM
Whi5-inducible 0 nM
Whi5-inducible 10 nM
Whi5-inducible 30 nM
b)
SCD
Mitochondrial diameter [nm]
replicate 1
replicate 2
replicate 3
WT 0 nM
WT 60 nM
WT 150 nM
Whi5-inducible 0 nM
Whi5-inducible 60 nM
Whi5-inducible 150 nM
c)
SCGE
Cell volume [fL]
0
100
200
300
400
500
600
700
800
replicate 1
replicate 2
replicate 3
WT 0 nM
WT 10 nM
WT 30 nM
Whi5-inducible 0 nM
Whi5-inducible 10 nM
Whi5-inducible 30 nM
d)
SCD
Cell volume [fL]
replicate 1
replicate 2
replicate 3
WT 0 nM
WT 60 nM
WT 150 nM
Whi5-inducible 0 nM
Whi5-inducible 60 nM
Whi5-inducible 150 nM
e)
SCGE
WT
β-estradiol inducible WHI5
0 nM β-estradiol
10 nM β-estradiol
30 nM β-estradiol
f)
SCD
WT
β-estradiol inducible WHI5
0 nM β-estradiol
60 nM β-estradiol
150 nM β-estradiol

**Supplementary Figure 4: Mitochondrial diameter does not depend on cell volume in yeast.** Wild-type (WT) and Whi5-inducible cells were grown in SCGE or SCD medium containing the indicated concentrations of β-estradiol, chemically fixed and analyzed by transmission electron microscopy. **a-d)** Quantification of cell volume and mitochondrial diameter corresponding to Fig. 2g. Box plots depict median (black line), 25th, 75th percentile (box). Whiskers indicate extreme values still within 1.5 interquartile ranges and outliers are depicted as single black points. Shown is the data from three independent experiments. In Fig. 2g, the mean of the means of all three replicates was plotted for the data for wild-type and Whi5-inducible cells grown without β-estradiol and Whi5-inducible cells grown in the presence of 10 nM and 30 nM (SCGE) or 60 nM and 150 nM β-estradiol (SCD). **a-b)** Mitochondrial diameter of cells grown in a) SCGE or b) SCD medium. For each sample, the diameter of 100 mitochondria was measured from electron micrographs. **c-d)** DIC images were used to calculate cell volume from cell segmentations performed with Cell-ACDC for cells grown in c) SCGE or d) SCD medium. The images were taken of live cells of the same cultures that were used for chemical fixation. **e-f)** Representative electron micrographs of mitochondria of wild-type and Whi5-inducible cells grown in e) SCGE or f) SCD medium containing the indicated concentrations of β-estradiol. Scale bars represent 500 nm.

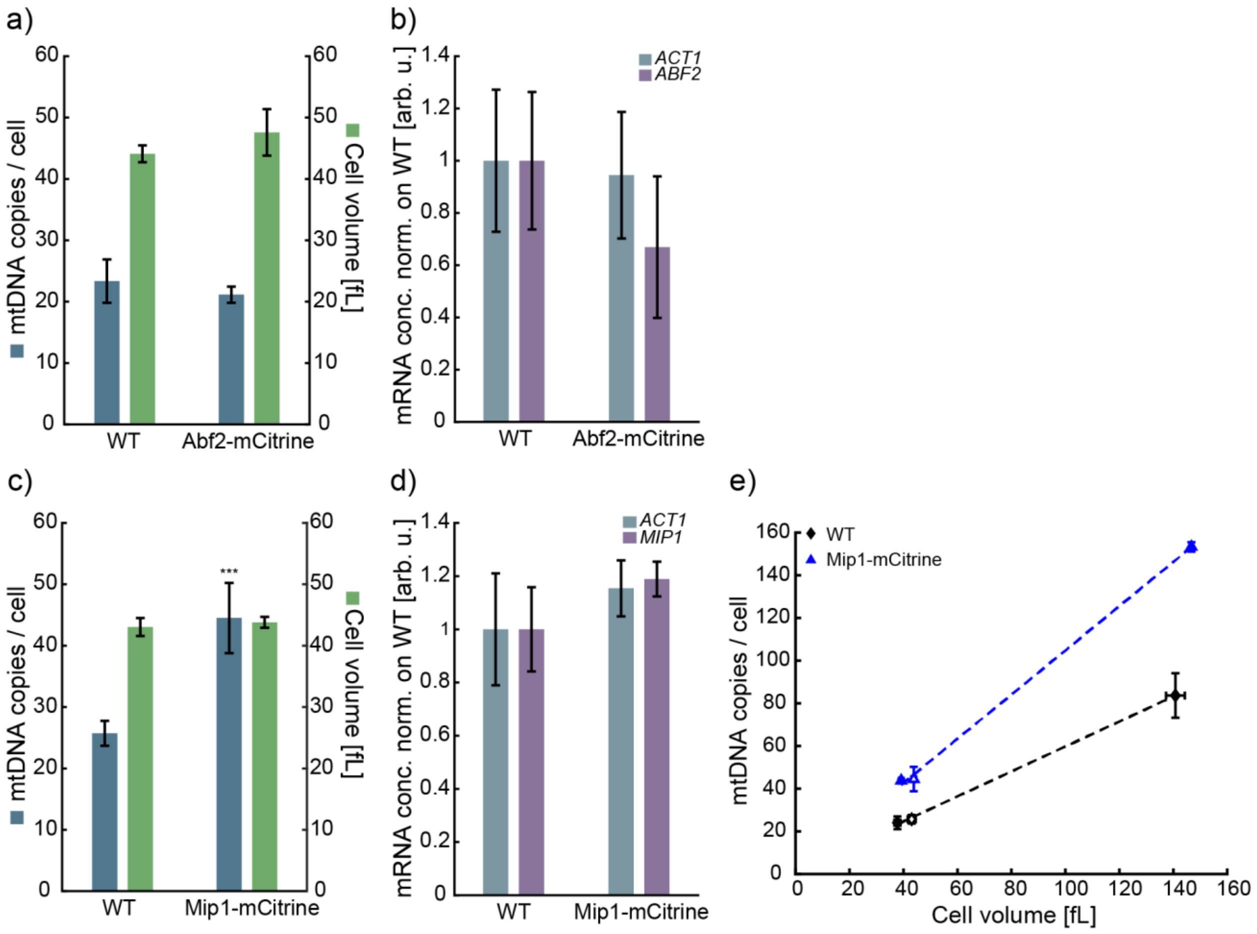


**Supplementary Figure 5: Abf2-mCitrine and Mip1-mCitrine are functional**. To measure mtDNA copy numbers, mean cell volume, and transcript levels in tagged strains, DNA-qPCRs, Coulter counter measurements and RT-qPCR were performed. **a-b)** We did not observe any significant changes of mtDNA copy number, cell volume (a) or *ACT1* and *ABF2* transcript levels (b) upon tagging Abf2 with mCitrine. **c-d)** Tagging Mip1 with mCitrine leads to a significant ($p < 0.001$) increase of mtDNA copy number by ~ 80%, while cell volume and transcript levels remain unchanged. **e)** To test if despite the change in mtDNA copy number, the cell-volume-dependent regulation of mtDNA is still intact, we tagged Mip1-mCitrine in a Whi5-inducible strain (filled triangles). We find that the Mip1-mCitrine strain still shows a cell volume-dependent increase of mtDNA.

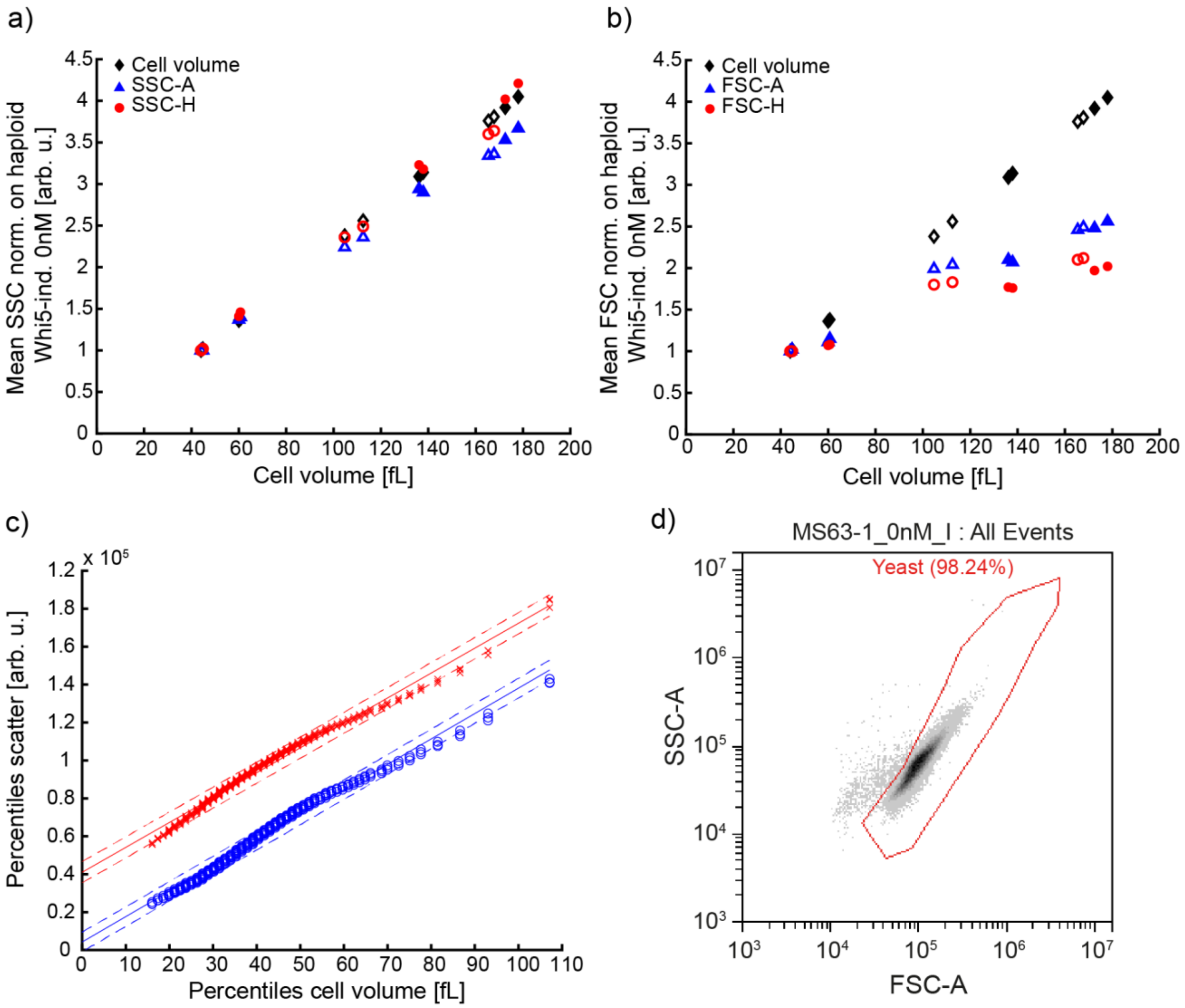


**Supplementary Figure 6: Comparison of flow cytometry side scatter and forward scatter measurements.** Whi5-inducible cell populations of haploid (open symbols) and diploid cells (filled symbols) were grown on SCGE with β-estradiol (haploids 0 nM, 10 nM, 30 nM; diploids 0 nM, 25 nM, 50 nM) and measured with a CytoFlex S Flow Cytometer (Beckman Coulter) in technical duplicates, recording SSC **(a)** and FSC **(b)**. Flow cytometry measurements were compared to mean cell volumes obtained from Coulter counter measurements. Blue triangles show area and red circles height measurements, both normalized on the Whi5-inducible strain grown with 0 nM β-estradiol. Normalized Coulter counter measurements are shown to guide the eye (black diamonds). We find that both side scatter measurements correlate well with Coulter counter cell volume measurements. **c)** We anticipate that SSC-A should be proportional to cell volume independent of cell-cycle stage. To test if this is the case, we compared the percentiles of scatter signal (SSC-A: blue circles, FSC-A: red crosses) with the percentiles of Coulter counter measurements for wild-type. Data of 3 biological replicates are shown. Lines show linear fits to the pooled data, with dashed lines showing 95% confidence prediction intervals. **d)** Example for gating strategy.

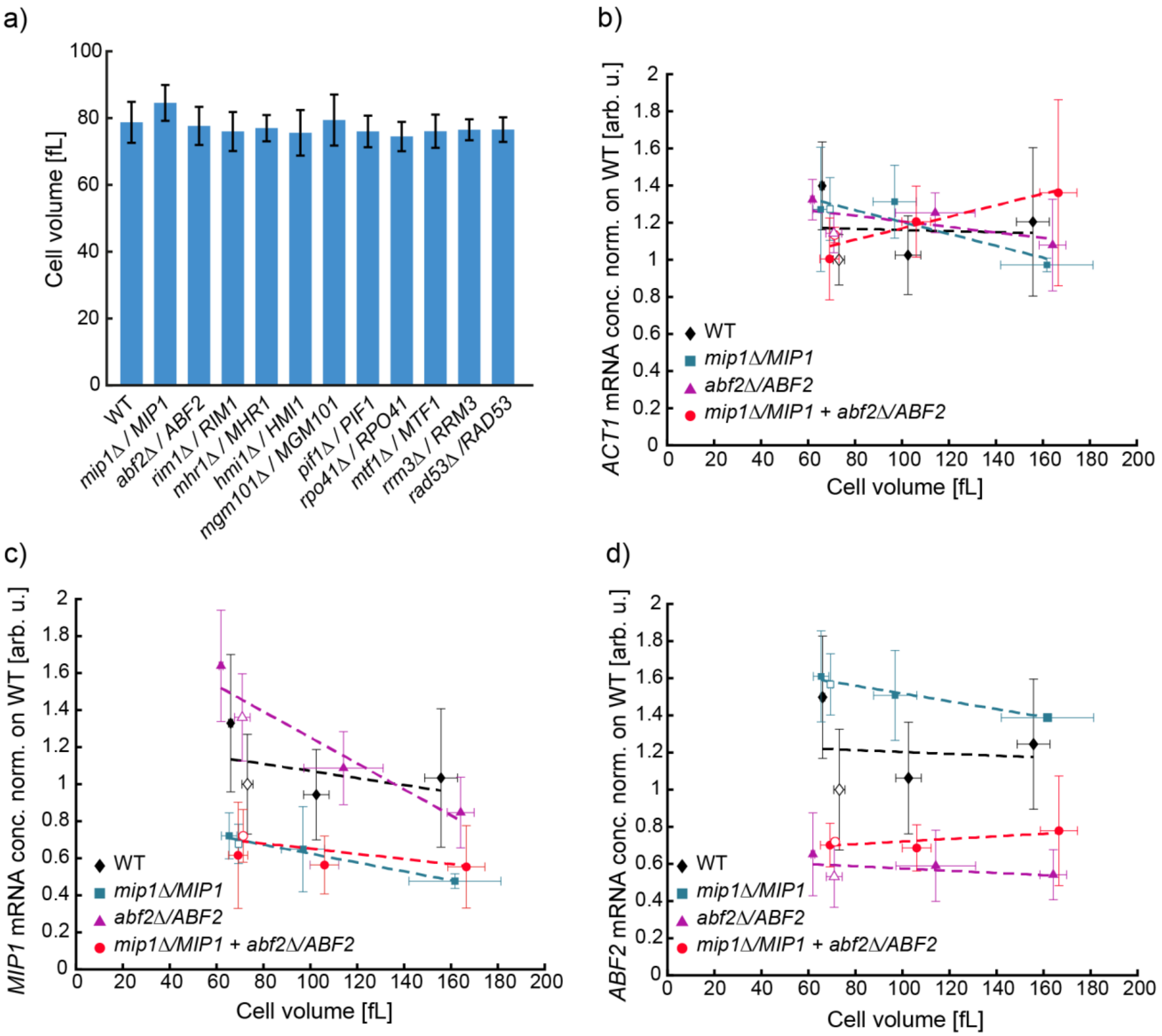


**Supplementary Figure 7. Hemizygous experiments for cell volume and mRNA levels. a)** Coulter counter measurements show no significant changes in cell volume for hemizygous mutants from Fig. 4b compared to wild-type. **b-d)** To verify that transcript levels are not dosage compensated in hemizygous *MIP1*, *ABF2* and double hemizygous strains, mRNA concentrations were measured by RT-qPCR for Whi5-inducible (0, 15, 60 nM β-estradiol; filled symbols) and non-inducible strains (open symbols) grown on SCGE, and normalized to the non-inducible wild-type (open diamond). **b)** *ACT1* concentrations stay similar for all strains. **c)** *MIP1* mRNA concentration is reduced by ~50% in single and double hemizygous strains. **d)** *ABF2* mRNA concentration is reduced by ~50% in single and double hemizygous strains.

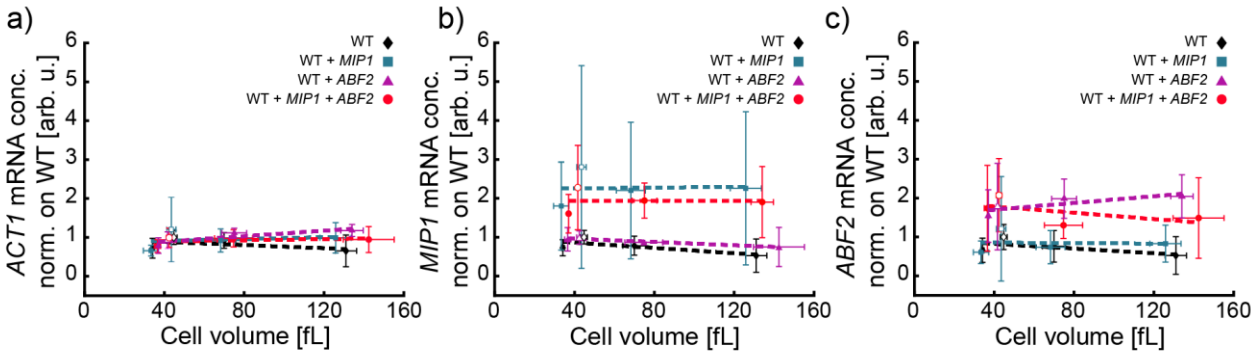


**Supplementary Figure 8: Multicopy strains tested for mRNA concentrations.** To verify that additional gene copies of *MIP1* and *ABF2* are expressed, mRNA concentrations were measured by RT-qPCR for Whi5-inducible (0, 10, 30 nM β-estradiol; filled symbols) and non-inducible strains (open symbols) grown on SCGE, and normalized on the non-inducible wild-type (open diamond). **a)** *ACT1* concentrations stay comparable for all strains. **b)** *MIP1* mRNA concentrations increase by ~ 100% in multicopy strains, where an additional *MIP1* allele was inserted. **c)** *ABF2* mRNA concentrations increase by ~ 100% in multicopy strains, where an additional *ABF2* allele was inserted.

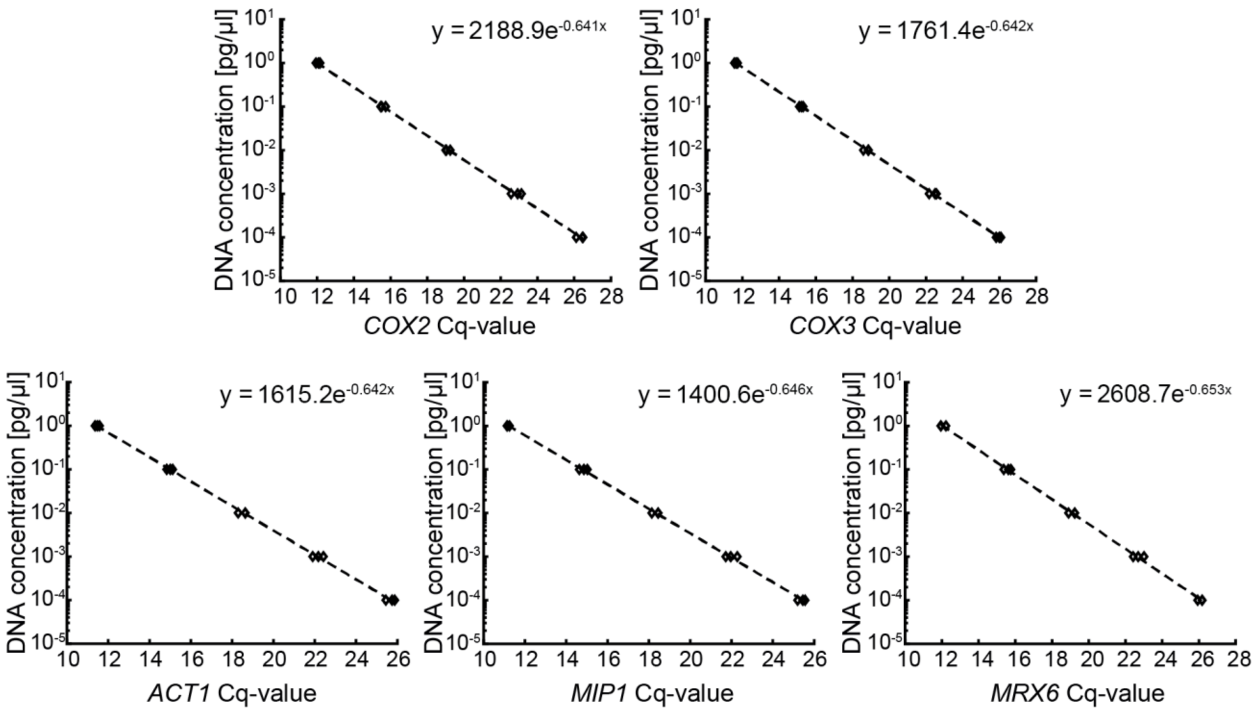


**Supplementary Figure 9: Standard curves of calibration standard.** A calibration standard was constructed by fusing the amplified sequences of *COX2*, *COX3*, *ACT1*, *MIP1* and *MRX6* to one template with PCR. Then a dilution series was prepared and qPCR was performed in three replicates for each primer pair. Results are shown in a semilog plot. By fitting the data, the equation of the standard curve was obtained and used to calculate concentrations of the corresponding genes for all DNA qPCR data.

**Table 1: Testing ASY39-1 ($\rho^0$ strain) for mtDNA by qPCR.** Raw Cq-values obtained for mitochondrial (*COX2, COX3*) and nuclear DNA (*ACT1*) sequences are shown for WT (MMY116-2c), haploid microscopy strain (ASY13-1) and $\rho^0$ strain (ASY39-1).

| Gene | WT Cq-value +/- StD | $\rho^+$ Cq-value +/- StD | $\rho^0$ Cq-value +/- StD |
|---|---|---|---|
| *COX2* | 18.96 +/-0.19 | 19.30 +/- 0.15 | Cq-values not measurable or higher than 36 |
| *COX3* | 18.53 +/- 0.30 | 18.73 +/- 0.15 | Cq-values not measurable or higher than 34 |
| *ACT1* | 22.83 +/- 0.33 | 22.70 +/- 0.33 | 21.84 +/- 0.26 |

**Table 2: Strains used in this study.**

| Name | Genotype | Description | Origin | Figure |
|---|---|---|---|---|
| ASY004-1 | *Mat α; ADE2, ABF2-linker-mCitrine-CglaTRP1* | ABF2-mCitrine in MMY116-2c | This study | 3a, Supp. Fig. 5a-b |
| ASY006-1 | *Mat α; ADE2, MIP1-linker-mCitrine- ADH1term-CglaTRP1* | MIP1-mCitrine in MMY116-2c | This study | 3a, Supp. Fig. 5c-e |
| ASY007-2 | *Mat a; ADE2, MIP1-linker-mCitrine:CglaTRP1, whi5Δ::kanMX6-LexApr-WHI5-ADH1term-LEU2, his3::LexA-ER-AD-TF-HIS3* | MIP1-mCitrine-Adh1term-CglaTRP1 (KCE001-2) in MS63-1 | This study | Supp. Fig. 5e |
| ASY013-1 | *Mat a; mt-LacO, ADE2, TRP1, whi5Δ::KlacURA3, LexApr-WHI5-ADH1term-LEU2, his3::LexA-ER-LBD-HIS3, Pcup1-Su9-2xNEON-LacI-Pgk1-Su9-mKate2-KanMX4* | Haploid microscopy strain, Whi5-inducible | This study | 2c-f; Supp. Fig. 3b-c; |
| ASY015-1 | *Mat α/a; mt-LacO, ADE2/ADE2, , Δwhi5::KlacURA3/WHI5, LexApr-WHI5-ADH1term-LEU2, his3::LexA-ER-LBD-HIS3, Pcup1-Su9-2xNEON-LacI-Pgk1-Su9-mKate2-KanMX4* | Diploid microscopy strain, Whi5-inducible | This study | 2b-d; Supp. Fig. 3a |
| ASY020-1 | *Mat α/a; ADE2/ADE2, URA3/ura3, leu2/LEU2* | Diploid WT | This study | 1b; 4b-c; 6a-b; Supp. Fig. 7, 8 |
| ASY023-1 | *Mat α/a; ADE2/ADE2, ura3-1/URA3, whi5Δ::kanMX6-LexApr-WHI5-ADH1term-LEU2/WHI5, his3::LexA-ER-AD-TF-HIS3/his3-11,15* | Whi5-inducible strain | This study | 1b; 6b |

| | | | | |
|---|---|---|---|---|
| ASY024-1 | *Mat α/a; ADE2/ADE2, leu2-3/LEU2, ura3-1/URA3, mip1::CgalTRP1/MIP1* | Hemizygous *MIP1* strain | This study | 4b-c; 6a-b; Supp. Fig. 7 |
| ASY025-1 | *Mat α/a; ADE2/ADE2, ura3-1/URA3, whi5Δ::kanMX6-LexApr-WHI5-ADH1term-LEU2/WHI5, his3::LexA-ER-AD-TF-HIS3/his3-11,15, mip1::CgalTRP1/MIP1* | Hemizygous *MIP1* strain, Whi5-inducible | This study | 6b; Supp. Fig. 7 b-d |
| ASY033-1 | *Mat α/a; ADE2/ADE2, leu2-3/LEU2, URA3/ura3-1, pif1::TRP1/PIF1* | Hemizygous *PIF1* strain | This study | 4b; Supp. Fig. 7a |
| ASY034-1 | *Mat α/a; ADE2/ADE2, leu2-3/LEU2, URA3/ura3-1, rad53::TRP1/RAD53* | Hemizygous *RAD53* strain | This study | 4b; Supp. Fig. 7a |
| ASY035-1 | *Mat α/a; ADE2/ADE2, leu2-3/LEU2, URA3/ura3-1, rrm3::TRP1/RRM3* | Hemizygous *RRM3* strain | This study | 4b; Supp. Fig. 7a |
| ASY039-1 | *Mat α/a; ADE2, ∆whi5::KlacURA3, hiWHI5:LEU2, LexA-ER-LBD:HIS3, Pcup1-Su9-2xNEON-LacI--Pgk1-Su9-mKate2:KanMX4, TRP1, mip1::clonNAT* | *mip1∆* microscopy strain | This study | 2e-f; Supp. Fig. 3b-c; Supp. Table 1 |
| ASY046-1 | *Mat α/a; ADE2/ADE2, leu2-3/LEU2, ura3-1/URA3, mip1::CgalTRP1/MIP1, abf2::clonNAT/ABF2* | Double hemizygous *MIP1 ABF2* strain | This study | 6a-b; Supp. Fig. 7 b-d |
| ASY049-2 | *Mat α/a; whi5Δ::kanMX6-LexApr-WHI5-ADH1term-LEU2, his3::LexA-ER-AD-TF-HIS3, mip1::CgalTRP1/MIP1, abf2::clonNAT/ABF2* | Double hemizygous *MIP1 ABF2* strain; Whi5-inducible | This study | 6b; Supp. Fig. 7 b-d |
| ASY051-2 | *Mat α; ADE2, ura3::URA3/ABF2* | Haploid WT with additional copy of *ABF2* | This study | 6d-e; Supp. Fig. 8 |
| ASY052-5 | *Mat a; ADE2, whi5Δ::kanMX6-LexApr-WHI5-ADH1term-LEU2, his3::LexA-ER-AD-TF-HIS3, ura3::URA3/ABF2* | Haploid Whi5-inducible strain with additional copy of *ABF2* | This study | 6e; Supp. Fig. 8 |
| ASY057-3 | *Mat α; ADE2, trp1::TRP1/MIP1* | Haploid WT with additional copy of *MIP1* | This study | 6d-e; Supp. Fig. 8 |
| ASY058-1 | *Mat a; ADE2, whi5Δ::kanMX6-LexApr-WHI5-ADH1term-LEU2, his3::LexA-ER-AD-TF-HIS3, trp1::TRP1/MIP1* | Haploid Whi5-inducible strain with additional copy of *MIP1* | This study | 6e; Supp. Fig. 8 |

| | | | | |
|---|---|---|---|---|
| ASY059-2 | *Mat α; ADE2, ura3::URA3/ABF2, trp1::TRP1/MIP1* | Haploid WT with additional *ABF2* and *MIP1* copies | This study | 6d-e; Supp. Fig. 8 |
| ASY060-1 | *Mat a; ADE2, whi5Δ::kanMX6-LexApr-WHI5-ADH1term-LEU2, his3::LexA-ER-AD-TF-HIS3, ura3::URA3/ABF2, trp1::TRP1/MIP1* | Haploid Whi5-inducible strain with additional copies of *ABF2* and *MIP1* | This study | 6e; Supp. Fig. 8 |
| JE611-c | *Mat α, cln1Δ, cln2Δ, cln3::leu2, lexOPr-Cln1-Leu2, ADE2, his3::cyc1-Pr-lexO TF-his3, TRP, URA* | Cln1-inducible strain | Jennifer Ewald, Skotheim lab | 1c; Supp. Fig. 2 |
| KSY244-1 | *Mat α/a; ADE2/ADE2, leu2-3/LEU2, URA3/ura3-1, abf2::TRP1/ABF2* | Hemizygous *ABF2* strain | This study | 4b,c; Supp. Fig. 7 |
| KSY245-1 | *Mat α/a; ADE2/ADE2, ura3-1/URA3, whi5Δ::kanMX6-LexApr-WHI5-ADH1term-LEU2/WHI5, his3::LexA-ER-AD-TF-HIS3/his3-11,15, abf2::CgalTRP1/ABF2* | Hemizygous *ABF2* strain; Whi5-inducible | This study | 6b; Supp. Fig. 7 b-d |
| MMY116-2c | *Mat α; ADE2* | Haploid WT strain | Skotheim lab stock | 1b, 2g-i; 6d-e; Supp. Fig 1, 4, 5a-d, 8 |
| MS63-1 | *Mat a; ADE2, whi5Δ::kanMX6-LexAprom-WHI5-ADH1term-LEU2, his3::LexA-ER-AD-TF-HIS3* | Haploid Whi5-inducible strain | Matthew Swaffer, Skotheim lab | 1b, 2g-i, 3a, 6e; Supp. Fig 1, 4, 5e, 6, 8 |
| KCY005-1 | Mat α/a; ADE2/ADE2, whi5Δ::CglaTRP1/whi5Δ::kanMX6-LexApromWHI5-ADH1term-LEU2, his3/his3::LexA-ERAD-TF-HIS3 | Diploid Whi5-inducible strain | Kora-Lee Claude, Schmoller lab | Supp. Fig. 6a-c |
| KSY246 | *Mat α/a; ADE2/ADE2, leu2-3/LEU2, URA3/ura3-1, hmi1::TRP1/HMI1* | Hemizygous *HMI1* strain | This study | 4b; Supp. Fig. 7a |
| KSY253 | *Mat α/a; ADE2/ADE2, leu2-3/LEU2, URA3/ura3-1, rpo41::TRP1/RPO41* | Hemizygous *RPO41* strain | This study | 4b; Supp. Fig. 7a |
| KSY254 | *Mat α/a; ADE2/ADE2, leu2-3/LEU2, URA3/ura3-1, mtf1::TRP1/MTF1* | Hemizygous *MTF1* strain | This study | 4b; Supp. Fig. 7a |
| KSY255 | *Mat α/a; ADE2/ADE2, leu2-3/LEU2, URA3/ura3-1, mhr1::TRP1/MHR1* | Hemizygous *MHR1* strain | This study | 4b; Supp. Fig. 7a |

| | | | | |
|---|---|---|---|---|
| KSY256 | *Mat α/a; ADE2/ADE2, leu2-3/LEU2, URA3/ura3-1, mgm101::TRP1/MGM101* | Hemizygous *MGM101* strain | This study | 4b; Supp. Fig. 7a |
| KSY257 | *Mat α/a; ADE2/ADE2, leu2-3/LEU2, URA3/ura3-1, rim1::TRP1/RIM1* | Hemizygous *RIM1* strain | This study | 4b; Supp. Fig. 7a |

**Table 3: qPCR primers used in this study**

| Gene | qPCR primer direction | qPCR primer sequence (5‘-3‘) |
|---|---|---|
| *ABF2* | forward | CCAACCTTACGTCCTGCTG |
| | reverse | CGTCAAACTCCTTCTTCGCC |
| *ACT1* | forward | CACCCTGTTCTTTTGACTGA |
| | reverse | CGTAGAAGGCTGGAACGTTG |
| *COX2* | forward | GTTGATGCTACTCCTGGTAGATT |
| | reverse | TTGCATGACCTGTCCCACAC |
| *COX3* | forward | TTGAAGCTGTACAACCTACC |
| | reverse | CCTGCGATTAAGGCATGATG |
| *MIP1* | forward | CCATCACAAGCAAGAACGGC |
| | reverse | GTCCCTTTCCAGCTCAACCA |
| *MRX6* | forward | CATCCGACGTGGTGCTCTTA |
| | reverse | TCTCATCTCTCCCTCCACCC |
| *MTF1* | forward | TTGCTAATGTGACGGGGGAG |
| | reverse | CTGTTGTGCTTGGCATCCAT |
| *PIM1* | forward | ACCCTACATTGGCGCTTTCA |
| | reverse | AGTGCCCGTCTTTTCGTCTT |
| *RDN18* | forward | AACTCACCAGGTCCAGACACAATAAGG |
| | reverse | AAGGTCTCGTTCGTTATCGCAATTAAGC |
| *RPO41* | forward | TCTGGGTAGAACACCGTGGA |
| | reverse | TTCGTCTTGTGCACCTGGAA |

**Table 4: Plasmids constructed in this study.** *MIP1* and *ABF2* promoters correspond to 1000 bp upstream of the start codon. *MIP1* terminator corresponds to 271 bp downstream of coding *MIP1*. *ABF2* terminator corresponds to 288 bp downstream of coding *ABF2*.

| Plasmid | Description |
|---|---|
| ASE001-5 | HO-homology-CuPr-SU9-2xmNeon-LacI |
| ASE002-2 | pRS404-Mip1Pr-Mip1-Mip1term |
| ASE003-1 | pRS406-Abf2Pr-Abf2-Abf2term |
